# The evolution of trait variance creates a tension between species diversity and functional diversity

**DOI:** 10.1101/2020.07.01.182261

**Authors:** György Barabás, Christine Parent, Andrew Kraemer, Frederik Van de Perre, Frederik De Laender

## Abstract

It seems intuitive that species diversity promotes functional diversity. For example, more plant species imply more diverse leaf chemistry and thus more kinds of food for herbivores. Here we argue that the evolution of functional trait variance challenges this view. We show that trait-based eco-evolutionary processes force species to evolve narrower trait breadths in tightly packed communities, in their effort to avoid competition with neighboring species. This effect is so strong as to reduce overall trait space coverage, overhauling the expected positive relationship between species- and functional diversity. Empirical data from Galápagos land snail communities proved consistent with this claim. As a consequence, trait data from species-poor communities may misjudge functional diversity in species-rich ones, and vice versa.

---

Functional traits are organismal traits impacting the ecological performance (fitness) of individuals [1,2]. They determine the way individuals interact within the community [3], and contribute to ecosystem functioning [4] and services [5]. The extent to which the available functional trait space is covered by the community is the community’s functional diversity [6–9], which is an important indicator of community structure and ecosystem health [8, 10]. (See the Supplementary Information [SI], Section 5 for various diversity measures.)

In a purely ecological setting, species diversity is a prime indicator of a community’s functional diversity: greater species diversity often implies greater functional diversity [11], even though the exact nature of this positive relationship might depend on factors such as latitude [12]. This positive association is a key component of theory in ecosystem and community ecology. For example, ecosystems with more species or greater evenness produce more biomass than monocultures, and do so more stably, because of a greater diversity of resource use strategies [13–15].

When also considering evolutionary processes however, the positive association between the two may be overhauled, as we argue here. We illustrate this point by comparing two communities, one with three and another with six extant species (Figure 1**AB**). Although the three-species community has lower species diversity, it is clearly more functionally diverse, as it covers a larger proportion of the trait axis. This is because its species have evolved much broader trait distributions. A similar example is shown in Figure 1**CD**, in a two-dimensional trait space.

**Figure 1:**
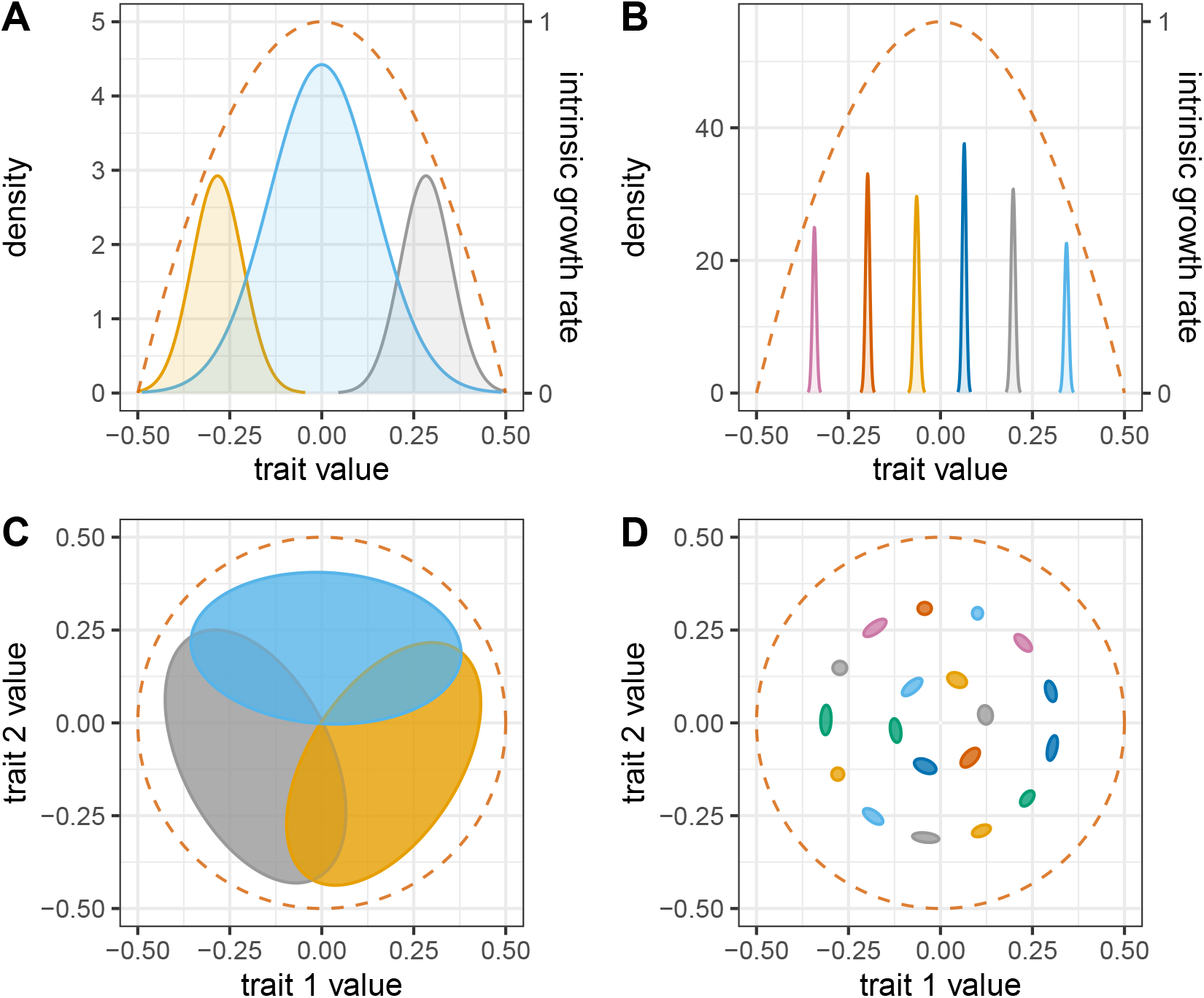
Greater species diversity does not necessarily translate to greater functional trait diversity. Panels show equilibrium states of simulated communities in one (**AB**) and two (**CD**) trait dimensions, with low (**AC**) and high (**BD**) final species diversity. In **A** and **B**, density is plotted along the ordinate and trait value along the abscissa. Trait distributions of different species are shown with different colors; the area under each curve is the total population density of the corresponding species. The dashed orange line shows what the rate of exponential growth of the given phenotype would be in the absence of competition (right-hand ordinate). The competition width (trait distance beyond which competition is significantly reduced between two phenotypes; SI, Section 6.1) is 0.15. In **C** and **D**, the axes correspond to the two traits, the contour lines are the 95% regions of trait variance, and the hue inside the contours is proportional to species’ relative densities. The dashed line encloses the region of trait space where intrinsic growth rates are positive; competition width is 0.2.

The examples of Figure 1 are not mere cartoons, but reflect typical outcomes of a general eco-evolutionary model, seeded with different initial numbers of species and returning communities that have reached an ecological and evolutionary equilibrium. This model for the first time integrates the evolution of an arbitrary number of traits [16, 17], species interactions [18], and does so in continuous time, which is computationally more efficient. Technically, the model is based on quantitative genetics: it tracks (i) the population density, and (ii) the mean and (iii) variance (or covariance matrix, in multiple trait dimensions) of each species’ trait distribution, assumed to be normal. These three quantities for each species fully characterize the state of the community, i.e., how individuals are distributed across trait space. Trait evolution and density change are driven by phenotypes experiencing differential intrinsic growth depending on their position in trait space, and by competition arising through the consumption of shared resources (SI, Sections 2-4). A major difference between the present model and existing approaches is the way phenotypic variance is handled. Most existing models keep this variance constant [19, 20], while our model considers trait variance evolution in a community context (SI, Sections 1-2).

The reason communities with more species diversity turn out less functionally diverse (Figure 1) is that they are more tightly packed with species. In such communities, species avoid competition by evolving very narrow trait variances, thus reducing their overlap with neighboring species. The stark reduction in species’ trait breadths can be understood by visualizing the fitness function over phenotypes (Figure 2). For low species diversity, the trait space has regions of both negative growth (in densely populated regions) and positive growth (in areas with lower prevalence of individuals, farther away from the densely occupied regions). The equilibrium trait breadth of each species then emerges as a balance between the trait-enhancing influence of positive growth regions and the trait-pruning influence of negative ones. But when species are tightly packed, the regions of positive growth shrink and ultimately disappear: the fitness function becomes negative everywhere except at species’ mean trait values, where it is zero. In time, this negative selection reduces the genetic variance of each species to zero, leaving only the environmental component of trait variation (SI, Section 1).

**Figure 2:**
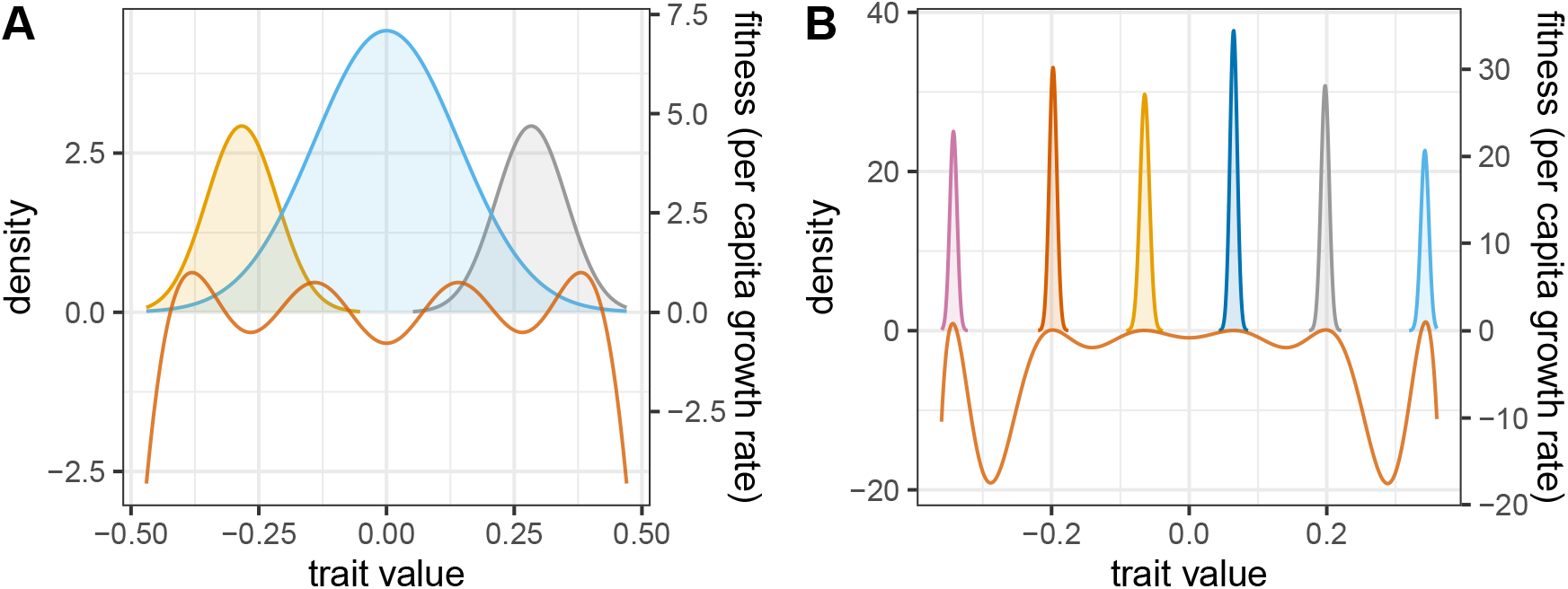
Graphical overview of why species evolve narrower trait breadths in more tightly packed communities. Shaded curves show species’ equilibrium trait distributions as in Figure 1; the orange line shows the per capita growth rate of the corresponding phenotype, independent of the species the phenotype belongs to. This fitness curve (right-hand ordinate) is given by Eq. 68 in the SI, after substituting in *b*(*z*) = 1 – 4*z*^2^. **A**: with fewer species, there are regions with both positive and negative growth along the trait axis, which will act to increase and decrease trait density in those regions, respectively. Thus, trait breadths evolve as a balance between the trait-enhancing vs. trait-pruning influence of these processes. **B**: in tightly packed communities, each phenotype experiences negative growth, except at species’ mean trait values, where growth is zero. This means that selection eventually removes all genetic variance. Any remaining trait variation is environmental, i.e., stemming from developmental noise and other sources unaffected by selection.

The outcome illustrated in Figures 1–2 is robust to changes in the model’s parameters and assumptions: functional diversity never increases monotonically with, and often declines with, species diversity (Figure 3). These results emerge from a numerical experimental setup whereby we vary the number of trait dimensions, the number of initial species, the amount of environmental trait covariance per species, the initial genetic covariance levels, the competition width, the shape of the intrinsic growth function, and the diversity metric used (SI, Section 6). Note that, in Figure 3, the effect of species diversity on functional diversity is weaker for larger environmental trait covariances. This stands to reason, as the total phenotypic covariance is the sum of the genetic and environmental covariances (SI, Section 1), of which only the former can evolve. If the environmental covariances are large, then no matter how tiny the genetic components evolve to be, the actual trait covariance (and therefore functional diversity) will still remain substantial.

**Figure 3:**
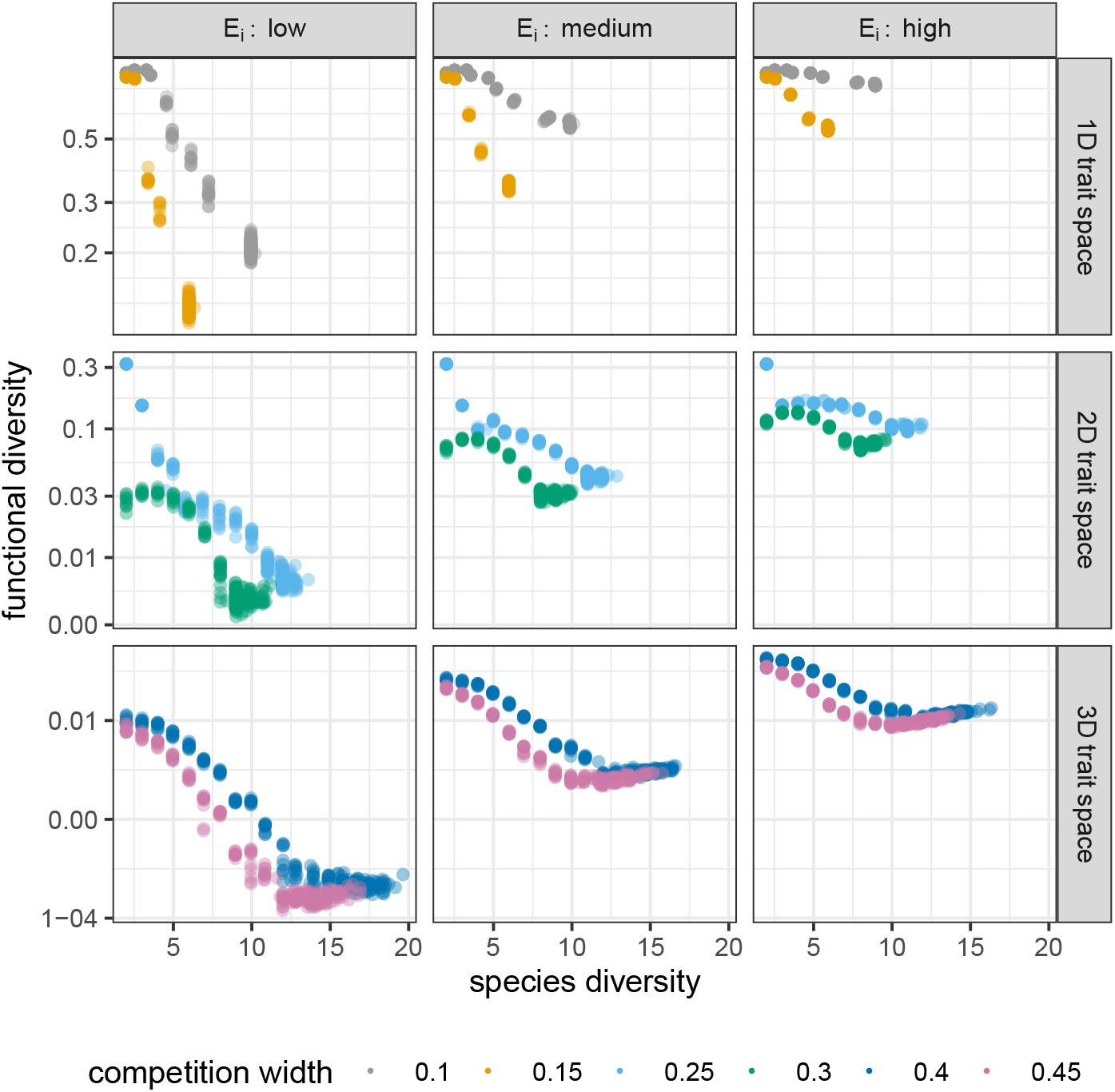
Functional diversity plotted against species diversity, for various dimensions of trait space (rows), levels of environmental trait variance (columns), and competition widths (colors). Intrinsic growth rates are given by *b*(*z*) = 1 – 16*z*^4^ (SI, Section 6.1). Functional diversity was obtained by binning each trait axis using a grid size of 1/100 in the [−1, 1] range (SI, Section 5; note the log scale along the ordinate). Points show results across ten replicates per parameterization. Since higher-dimensional trait spaces have more room and therefore can, all other things being equal, harbor more species, the competition widths were chosen larger for higher-dimensional trait spaces to create comparable species diversities for different trait dimensionality.

The model does have some limitations, nuancing the interpretation of the numerical results. First, there is no upper limit to genetic covariances, which is an artifact of having assumed large populations and infinitely many alleles per locus, instead of modeling individuals and their alleles explicitly (SI, Section 1). This means that the model may predict increases in trait variance which would not be possible with realistic levels of standing genetic variation. Second, since trait distributions are always normal and therefore unimodal, the model cannot produce speciation (which requires a multimodal trait distribution). That is, in our model, only the parameters of this distribution can evolve (mean and variance), not the type of distribution. Disruptive selection within a single species will therefore increase this species’ trait variance without ever splitting into two species. Including speciation into our model would by definition increase species diversity, while possibly reducing trait diversity, thus tempering any negative relationship between the two. In sum, the model almost certainly overestimates the reduction in functional diversity, and it is therefore safer to say that we expect the lack of a positive relationship, rather than a necessarily negative one, between species- and functional diversity.

While this mechanism, over evolutionary time, should leave its mark on any community conforming to its assumptions, directly quantifying its effects is hampered by the fact that diversity patterns in natural communities are typically driven by a suite of other local and regional processes. One possible approach is to identify a suitable “natural evolution experiment”: a set of isolated subcommunities that have near-identical biotic and abiotic environmental conditions, but have evolved independently for a long time, and also vary in the species diversity of a focal functional group. Then, if individuals vary in some functional traits which also mediate competition, and these traits have been recorded at the individual level, we can evaluate whether relatively more species-diverse subcommunities tend to have relatively lower functional diversity.

We have found only one natural evolution experiment which conforms to the assumptions of our model and for which the necessary data are available. It is a dataset of endemic land snails on the Galápagos Islands (SI, Section 7), from the genus *Naesiotus*. Previous work has found that intraspecific variation in the land snails’ shell morphology is positively correlated with habitat heterogeneity and negatively correlated with the number of co-occurring congener species. These together suggest that snails compete for available niche space, and that shell morphology reflects adaptedness to those ecological opportunities [21]. Displaying the distribution of individuals in each subcommunity reveals that species segregate in the two-dimensional trait space spanned by shell centroid size and shell shape (measured by the first PC axis explaining over 80% of shape variation), further supporting the idea that shell morphology, quantified by these two traits, mediates competition (Figure S10). Community age varies from 60 thousand to over three million years across the islands [21, 22], which therefore have had sufficient time for shell morphology to evolve. Some unrelated species occurring on different islands have highly similar shell morphotypes [23], strongly implying that evolution has converged on similar solutions, and therefore that the species form evolutionarily stable communities.

There are thirteen islands in the dataset, most of which possess both arid and humid habitat zones, with the distribution ranges of snail species never overlapping across them. Thus, their species do not have the opportunity to interact with one another, and so the compositions of the humid and arid zones form effectively separate communities. We used the shell morphology of individual snails to compute functional diversity for each subcommunity (i.e., island-habitat zone combination). To appropriately evaluate species diversity, we had to account for the (sometimes vastly) varying habitat heterogeneity across islands. Snails tend to partition habitat based on structures and surfaces available (e.g., under rocks or logs, low on tree trunks, on small low vegetation). Therefore, although the snails are not host-specialists, they have clear microhabitat preferences (C. Parent, personal observation), and ecological opportunities are thought to roughly scale with the number of available host plant species per community [21]. Finally, one limitation of the dataset is that the recorded relative species abundances do not necessarily reflect the actual ones on the islands, which would be needed for diversity calculations. We therefore also tested the sensitivity of the empirical relationship between species- and functional diversity to randomizing these relative abundances.

The data do not support a positive relationship between functional diversity and species diversity *per host plant species*, and even suggest a negative one (Figure 4). Even if one does not correct for the number of host plant species and considers raw species diversity, no positive relationship is found (SI, Section 7.2). Randomizing relative abundances does not change this result, showing that this limitation of the dataset likely does not affect our conclusions (SI, Section 7.2).

**Figure 4:**
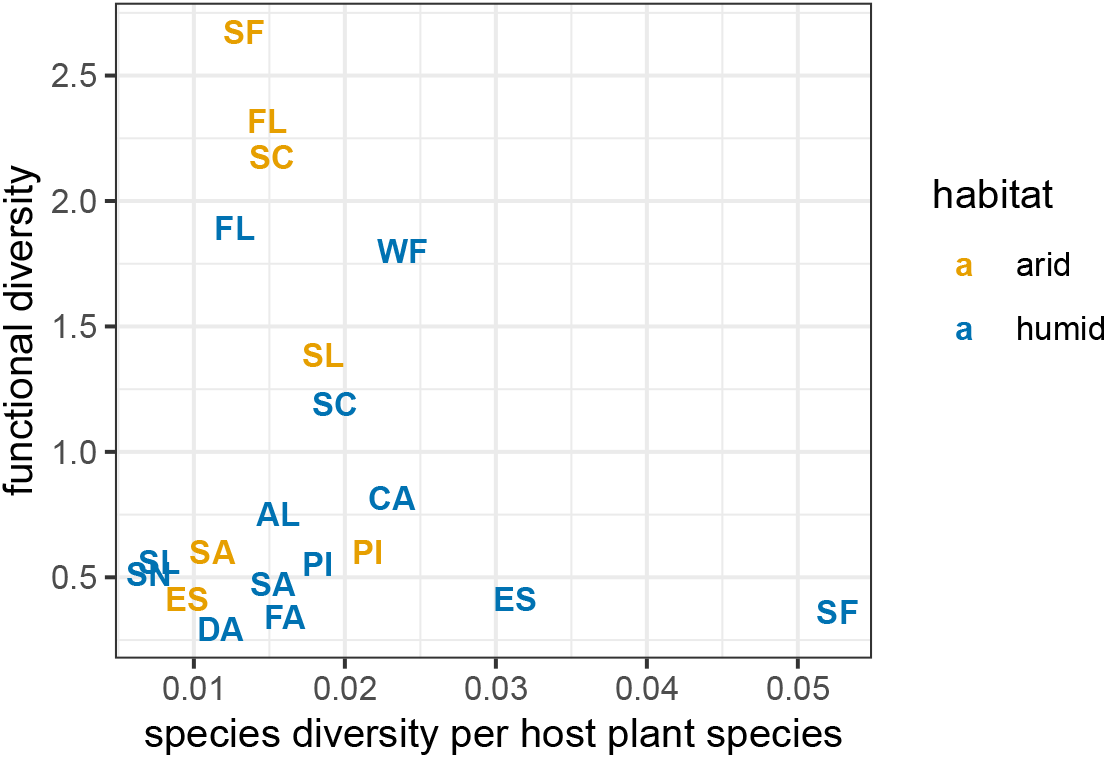
Functional diversity plotted against species diversity, for the land snail communities on the Galápagos Islands. Species diversity is normalized by the number of host plant species in each subcommunity, to get a better index of species diversity relative to the number of available ecological opportunities. Labels are island name abbreviations: Alcedo Volcano (AL), Cerro Azul Volcano (CA), Darwin Volcano (DA), Espanola (ES), Fernandina (FA), Floreana (FL), Pinzon (PI), Santiago (SA), Santa Cruz (SC), Santa Fe (SF), San Cristobal (SL), Sierra Negra Volcano (SN), Wolf Volcano (WF). Colors show communities in the arid (yellow) and humid (blue) zones of the islands, which form independent subcommunities. The data do not support a positive relationship between species- and functional diversity, and even offer evidence for a negative one. (A linear regression has slope −11.4± 17.5 with *p* = 0.53; however, since the data are heteroscedastic and the expected patterns are not linear to begin with, any such statistic should be treated as just an illustration.) This trend is retained after randomizing the number of sampled individuals (SI, Section 7.2), showing that the results are robust against measurement error in relative abundances and also lending further support to the relationship being actually negative.

Our main result that the relationship between species- and functional diversity tends to be neutral or negative rather than positive and monotonic has implications for the study of ecosystem functioning. However, these implications are not necessarily immediate or obvious. Unless there is evidence that a given function is directly proportional to functional diversity (in which case lower functional diversity by definition maps to lower function), one must first determine how that function depends on available traits. For instance, our model can be used to explore two commonly studied ecosystem functions: resource use and biomass production [14, 15]. It turns out that even with trait variance evolution reducing functional diversity, species diversity continues to promote these two functions (SI, Section 6.3)—even though the degree of promotion could have been greater if the species had not adapted their trait variances to the presence of competitors. This is because the low functional diversity found in communities with high species diversity is still sufficiently high to achieve positive complementarity [24].

In conclusion, we have found both theoretical and empirical evidence that evolutionary processes can impede a positive monotonic relationship between species- and functional diversity, and may even turn it into a negative one. The reason is that species evolve narrow trait breadths in species-rich communities to avoid competition with neighbors. That intraspecific trait variation decreases with species richness has been argued before [25–27]. Our results, for the first time, show that evolution can make this reduction so strong as to curtail the overall coverage of trait space. This reinforces the message that functional diversity should be measured at the individual level instead of aggregating traits into species averages [25–29]. Crucially, our results also highlight a new reason why doing so is important: species’ contributions to functional diversity obtained in species-poor communities, even if individual-based, can overestimate functional diversity in species-rich communities.

## Acknowledgments

We thank G. Meszéna, J. Årevall, C. de Mazancourt, and M. Loreau for discussions, and F. Barraquand for helpful comments on our manuscript draft. Snail specimens were collected under permits from the Charles Darwin Foundation and the Galápagos National Park Directorate who also provided logistical help that made this work possible (CDF: #044-06, GNPD: #PC-45-14, PC-52-15, PC-52-16). We thank C. Sevilla, W. Cabrera, N. Castillo, T. de Roy, N. Carter, C. Philip, C. Philson, Z. Root, Y. Roell, and B. Miller for invaluable logistic, field, and laboratory assistance.

## Funding

GB was funded by the Swedish Research Council (grant VR 2017-05245). Galápagos field work was supported by grants from the National Institute of General Medical Sciences of the National Institutes of Health (IDeA #P30 GM103324), the National Science Foundation (#1523540 to ACK and #1751157 to CEP), the National Geographic Society, the American Malacological Society, the Western Society of Malacology, the Conchologists of America, and the Systematics Research Fund to CEP. FVdP was supported by a PhD fellowship from the Research Foundation Flanders. FDL supported from grants of the University of Namur (FSR Impulsionnel 48454E1); the Fund for Scientific Research, FNRS (PDR T.0048.16); and the ARC grant DIVERCE, a concerted research action from the special research fund.

## Author contributions

GB and FDL conceived of the study, ran simulations, and analyzed data; GB developed the theory and wrote the manuscript and Supplementary Information; CP & AK provided snail data. All authors contributed to the final form of the manuscript and helped with data analysis.

## Competing interests

There are no competing interests.

## Data availability

Data on the Galápagos land snail communities, along with code to replicate our analyses, can be accessed from www.github.com/dysordys/phenotypediv.

## Notes

https://www.github.com/dysordys/phenotypediv

### Competing Interest Statement

The authors have declared no competing interest.

