## Supplementary Information for "The evolution of trait variance creates a tension between species diversity and functional diversity"

György Barabás*, Christine Parent, Andrew Kraemer,
Frederik Van de Perre & Frederik De Laender

#### 1 Quantitative genetics of multidimensional traits

##### 1.1 Basic relationships

Let an individual's multidimensional phenotype $\boldsymbol{p}$ be an $L$-component vector, each entry holding the value of the corresponding character for that individual. We assume random mating, equal sex ratios, and purely additive genetic variation without genotype-environment interactions, dominance, epistasis, or pleiotropy. The multivariate normal distribution $\mathcal{N}(\boldsymbol{z}; \boldsymbol{\mu}, \boldsymbol{P})$ with mean $\boldsymbol{\mu}$, covariance matrix $\boldsymbol{P}$, and independent variable $\boldsymbol{z}$ is written

$$\mathcal{N}(\boldsymbol{z}; \boldsymbol{\mu}, \boldsymbol{P}) = \sqrt{\frac{1}{(2\pi)^L \det(\boldsymbol{P})}} \exp\left(-\frac{1}{2}(\boldsymbol{z}-\boldsymbol{\mu})\boldsymbol{P}^{-1}(\boldsymbol{z}-\boldsymbol{\mu})\right). \tag{1}$$

The phenotype $\boldsymbol{p}$ is composed of independent genetic and environmental factors, each of which is assumed to be sampled from an $L$-normal distribution:

$$\boldsymbol{p} = \boldsymbol{m} + \boldsymbol{f} + \boldsymbol{e}, \tag{2}$$

where $\boldsymbol{m}$ and $\boldsymbol{f}$ are the genetic contribution of the individual's mother and father, and $\boldsymbol{e}$ is the environmental contribution. While this description is purely statistical, it helps to think of the genetic component in the following way: each entry of $\boldsymbol{m}$ and $\boldsymbol{f}$ represents a single locus involved in coding for the corresponding trait, each locus having very many possible alleles.

The variance-covariance matrix $\mathrm{cov}(\boldsymbol{p})$ of the phenotype vector is calculated as

$$\begin{aligned}\mathrm{cov}(\boldsymbol{p}) &= \mathrm{cov}(\boldsymbol{m}+\boldsymbol{f}+\boldsymbol{e}) \\ &= \mathrm{cov}(\boldsymbol{m}) + \mathrm{cov}(\boldsymbol{f}) + \mathrm{cov}(\boldsymbol{e}) + 2\mathrm{cov}(\boldsymbol{m},\boldsymbol{f}) + 2\mathrm{cov}(\boldsymbol{m},\boldsymbol{e}) + 2\mathrm{cov}(\boldsymbol{f},\boldsymbol{e}),\end{aligned} \tag{3}$$

where $\mathrm{cov}(\boldsymbol{x},\boldsymbol{y})$ is the cross-covariance between two vectors $\boldsymbol{x}$ and $\boldsymbol{y}$. Since $\boldsymbol{m}$ and $\boldsymbol{f}$ stem from independently derived alleles, $\mathrm{cov}(\boldsymbol{m},\boldsymbol{f}) = 0$. Similarly, due to the lack of genotype-environment interactions, the other two cross-covariance terms are zero as well. This leaves

$$\mathrm{cov}(\boldsymbol{p}) = \mathrm{cov}(\boldsymbol{m}) + \mathrm{cov}(\boldsymbol{f}) + \mathrm{cov}(\boldsymbol{e}). \tag{4}$$

We will denote $\mathrm{cov}(\boldsymbol{m}) = \mathrm{cov}(\boldsymbol{f})$ by $\boldsymbol{G}/2$, where $\boldsymbol{G}$ is the additive genetic variance-covariance matrix. Similarly, we write $\boldsymbol{E} = \mathrm{cov}(\boldsymbol{e})$ and $\boldsymbol{P} = \mathrm{cov}(\boldsymbol{p})$. Eq. 4 then reads

$$\boldsymbol{P} = \boldsymbol{G} + \boldsymbol{E}, \tag{5}$$

with $\boldsymbol{P}$ being the sum of the additive genetic and environmental covariance matrices.

Next, we determine the covariance between a single parent and her offspring. Let their phenotypes be

$$\boldsymbol{p}_p = \boldsymbol{m}_p + \boldsymbol{f}_p + \boldsymbol{e}_p, \qquad \boldsymbol{p} = \boldsymbol{m}_p + \boldsymbol{f} + \boldsymbol{e}, \tag{6}$$

where the subscript $p$ denotes parental quantities. The maternal allele of the offspring, for each trait, is received from its mother, therefore $\boldsymbol{m} = \boldsymbol{m}_p$ was used above. The cross-covariance reads

$$\begin{aligned}\mathrm{cov}(\boldsymbol{p}_p, \boldsymbol{p}) = \mathrm{cov}(\boldsymbol{m}_p + \boldsymbol{f}_p + \boldsymbol{e}_p, \boldsymbol{m}_p + \boldsymbol{f} + \boldsymbol{e}) = \mathrm{cov}(\boldsymbol{m}_p) + \mathrm{cov}(\boldsymbol{m}_p, \boldsymbol{f}) + \mathrm{cov}(\boldsymbol{f}_p, \boldsymbol{m}_p) \\ + \mathrm{cov}(\boldsymbol{f}_p, \boldsymbol{f}) + \mathrm{cov}(\boldsymbol{m}_p, \boldsymbol{e}) + \mathrm{cov}(\boldsymbol{f}_p, \boldsymbol{e}) + \mathrm{cov}(\boldsymbol{e}_p, \boldsymbol{m}_p) + \mathrm{cov}(\boldsymbol{e}_p, \boldsymbol{f}) + \mathrm{cov}(\boldsymbol{e}_p, \boldsymbol{e}).\end{aligned} \tag{7}$$

The first term is $\mathrm{cov}(\boldsymbol{m}_p) = \mathrm{cov}(\boldsymbol{m}) = \boldsymbol{G}/2$. Terms 2-4 are zero because they are cross-covariances between independently derived alleles. Terms 5-8 are also zero due to the absence of genotype-environment interactions. Finally, we assume that the correlation between parent and offspring environments is negligible, setting $\mathrm{cov}(\boldsymbol{e}_p, \boldsymbol{e})$ to zero. In the end, we have

$$\mathrm{cov}(\boldsymbol{p}_p, \boldsymbol{p}) = \frac{1}{2}\boldsymbol{G}. \tag{8}$$

This turns out to be the same as the midparent-offspring covariance. The midparent phenotype reads $\boldsymbol{p}_m = (\boldsymbol{p}_1 + \boldsymbol{p}_2)/2$, from which

$$\mathrm{cov}(\boldsymbol{p}_m, \boldsymbol{p}) = \mathrm{cov}\left(\frac{\boldsymbol{p}_1 + \boldsymbol{p}_2}{2}, \boldsymbol{p}\right) = \frac{1}{2}\Big[\underbrace{\mathrm{cov}(\boldsymbol{p}_1, \boldsymbol{p})}_{\boldsymbol{G}/2} + \underbrace{\mathrm{cov}(\boldsymbol{p}_2, \boldsymbol{p})}_{\boldsymbol{G}/2}\Big] = \frac{1}{2}\boldsymbol{G}. \tag{9}$$

The variance-covariance matrix of midparent phenotype reads

$$\mathrm{cov}(\boldsymbol{p}_m) = \mathrm{cov}\left(\frac{\boldsymbol{p}_1 + \boldsymbol{p}_2}{2}\right) = \frac{1}{4}\underbrace{\mathrm{cov}(\boldsymbol{p}_1)}_{\boldsymbol{P}} + \frac{1}{4}\underbrace{\mathrm{cov}(\boldsymbol{p}_2)}_{\boldsymbol{P}} + \frac{1}{2}\underbrace{\mathrm{cov}(\boldsymbol{p}_1, \boldsymbol{p}_2)}_{0} = \frac{1}{2}\boldsymbol{P}. \tag{10}$$

#### 1.2 Midparent-offspring regression without selection

To perform the regression of offspring on midparents, we construct a $2L$-dimensional multi-normal distribution whose first $L$ dimensions correspond to midparents, and the second $L$ to offspring. Its mean $\boldsymbol{\mu}_{p_m,p}$ and covariance matrix $\boldsymbol{\Sigma}_{p_m,p}$ can be written in block-matrix form:

$$\boldsymbol{\mu}_{p_m,p} = \begin{pmatrix}\boldsymbol{\mu} \\ \boldsymbol{\mu}\end{pmatrix}, \qquad \boldsymbol{\Sigma}_{p_m,p} = \begin{pmatrix}\mathrm{cov}(\boldsymbol{p}_m) & \mathrm{cov}(\boldsymbol{p}_m, \boldsymbol{p}) \\ \mathrm{cov}(\boldsymbol{p}, \boldsymbol{p}_m) & \mathrm{cov}(\boldsymbol{p})\end{pmatrix}, \tag{11}$$

where the mean of midparents and offspring are equal due to the lack of selection (note also that the midparent and individual parental mean phenotypes are equal). Using $\boldsymbol{P} = \mathrm{cov}(\boldsymbol{p})$ and Eqs. 9-10, the covariance terms above can be written as

$$\boldsymbol{\Sigma}_{p_m,p} = \begin{pmatrix}\boldsymbol{P}/2 & \boldsymbol{G}/2 \\ \boldsymbol{G}/2 & \boldsymbol{P}\end{pmatrix}. \tag{12}$$

Using standard multivariate regression (Eaton 1983, pp. 116-117), the expected offspring phenotype $\boldsymbol{\mu}'$ given a particular midparent phenotype $\boldsymbol{x}$ is

$$\boldsymbol{\mu}'(\boldsymbol{x}) = \mathrm{cov}(\boldsymbol{p}_m, \boldsymbol{p})\mathrm{cov}(\boldsymbol{p}_m)^{-1}(\boldsymbol{x} - \boldsymbol{\mu}) + \boldsymbol{\mu} = \boldsymbol{G}\boldsymbol{P}^{-1}(\boldsymbol{x} - \boldsymbol{\mu}) + \boldsymbol{\mu}, \tag{13}$$

and the covariance of offspring (also known as the the segregation variance) reads

$$\boldsymbol{S} = \mathrm{cov}(\boldsymbol{p}) - \mathrm{cov}(\boldsymbol{p}_m, \boldsymbol{p})\mathrm{cov}(\boldsymbol{p}_m)^{-1}\mathrm{cov}(\boldsymbol{p}_m, \boldsymbol{p}) = \boldsymbol{P} - \frac{1}{2}\boldsymbol{G}\boldsymbol{P}^{-1}\boldsymbol{G}. \tag{14}$$

The total phenotypic covariance of offspring is composed of two parts. First, the covariance matrix of $\boldsymbol{\mu}'$. Second, since, $\boldsymbol{\mu}'$ is the expected offspring phenotype given a midparent phenotype, one has to add the segregation variance $\boldsymbol{S}$ to this result. The two are indeed additive without cross-covariance terms, since a regression is always uncorrelated with its residuals. This means that the total phenotypic covariance of offspring can be written

$$\boldsymbol{P}' = \mathrm{cov}(\boldsymbol{\mu}') + \boldsymbol{S} = \mathrm{cov}\Big(\boldsymbol{G}\boldsymbol{P}^{-1}(\boldsymbol{x} - \boldsymbol{\mu}) + \boldsymbol{\mu}\Big) + \boldsymbol{P} - \frac{1}{2}\boldsymbol{G}\boldsymbol{P}^{-1}\boldsymbol{G}. \tag{15}$$

Using the identity $\mathrm{cov}(\boldsymbol{A}\boldsymbol{x} + \boldsymbol{a}) = \boldsymbol{A}\mathrm{cov}(\boldsymbol{x})\boldsymbol{A}^{\mathrm{T}}$ (where $\boldsymbol{A}$ is a constant matrix, $\boldsymbol{A}^{\mathrm{T}}$ its transpose, $\boldsymbol{a}$ a constant vector, and $\boldsymbol{x}$ the random vector), we get

$$\begin{aligned} \boldsymbol{P}' &= \mathrm{cov}\Big(\boldsymbol{G}\boldsymbol{P}^{-1}\boldsymbol{x} - \big(\boldsymbol{G}\boldsymbol{P}^{-1}\boldsymbol{\mu} + \boldsymbol{\mu}\big)\Big) + \boldsymbol{P} - \frac{1}{2}\boldsymbol{G}\boldsymbol{P}^{-1}\boldsymbol{G} \\ &= \boldsymbol{G}\boldsymbol{P}^{-1}\mathrm{cov}(\boldsymbol{x})\boldsymbol{P}^{-1}\boldsymbol{G} + \boldsymbol{P} - \frac{1}{2}\boldsymbol{G}\boldsymbol{P}^{-1}\boldsymbol{G}, \end{aligned} \tag{16}$$

where we used the fact that $(\boldsymbol{G}\boldsymbol{P}^{-1})^{\mathrm{T}} = (\boldsymbol{P}^{-1})^{\mathrm{T}}\boldsymbol{G}^{\mathrm{T}} = \boldsymbol{P}^{-1}\boldsymbol{G}$, since both $\boldsymbol{G}$ and $\boldsymbol{P}$ are covariance matrices and thus symmetric. By Eq. 5, $\boldsymbol{P} = \boldsymbol{G} + \boldsymbol{E}$ and $\boldsymbol{P}' = \boldsymbol{G}' + \boldsymbol{E}$ (assuming a constant environmental covariance). Inserting the identity matrix $\boldsymbol{P}\boldsymbol{P}^{-1}$ into the last term:

$$\boldsymbol{P}' = \boldsymbol{G}' + \boldsymbol{E} = \boldsymbol{G}\boldsymbol{P}^{-1}\mathrm{cov}(\boldsymbol{x})\boldsymbol{P}^{-1}\boldsymbol{G} + \boldsymbol{G} + \boldsymbol{E} - \frac{1}{2}(\boldsymbol{G}\boldsymbol{P}^{-1})\boldsymbol{P}(\boldsymbol{P}^{-1}\boldsymbol{G}), \tag{17}$$

or

$$\boldsymbol{G}' = \boldsymbol{G}\boldsymbol{P}^{-1}\left(\mathrm{cov}(\boldsymbol{x}) - \frac{1}{2}\boldsymbol{P}\right)\boldsymbol{P}^{-1}\boldsymbol{G} + \boldsymbol{G}. \tag{18}$$

Eqs. 13 and 18 can also be written more transparently in terms of the difference between values at the two generations, $\Delta\boldsymbol{\mu} = \boldsymbol{\mu}' - \boldsymbol{\mu}$ and $\Delta\boldsymbol{G} = \boldsymbol{G}' - \boldsymbol{G}$:

$$\Delta\boldsymbol{\mu} = \boldsymbol{G}\boldsymbol{P}^{-1}(\boldsymbol{x} - \boldsymbol{\mu}), \tag{19}$$

$$\Delta\boldsymbol{G} = \boldsymbol{G}\boldsymbol{P}^{-1}\left(\mathrm{cov}(\boldsymbol{x}) - \frac{1}{2}\boldsymbol{P}\right)\boldsymbol{P}^{-1}\boldsymbol{G}. \tag{20}$$

#### 1.3 Selection on the parents

The regression results above hold in the absence of selection. We now consider what happens when the mean parental phenotype changes from the original $\boldsymbol{\mu}$ to a new $\tilde{\boldsymbol{\mu}}$ due to selection.

Eq. 19 gives the expected phenotype change as a function of the midparent phenotype $\boldsymbol{x}$. The mean midparent phenotype is in turn $\boldsymbol{\mu}$ without selection (which is also equal to the mean parental phenotype). Without selection, substituting $\boldsymbol{x} = \boldsymbol{\mu}$ in Eq. 19 predicts no expected phenotype change. With selection however, $\boldsymbol{x}$ should be replaced by $\tilde{\boldsymbol{\mu}}$. Then Eq. 19 reads

$$\Delta\boldsymbol{\mu} = \boldsymbol{G}\boldsymbol{P}^{-1}(\tilde{\boldsymbol{\mu}} - \boldsymbol{\mu}), \tag{21}$$

which is the breeder's equation for multiple characters. The matrix $\boldsymbol{G}\boldsymbol{P}^{-1}$ is the multidimensional analogue of the heritability $h^2 = G/P$. Note that quantities in Eq. 21 now refer to parents instead of midparents.

The change in additive genetic covariance is given by Eq. 20. Again, without selection this leads to no change, because

$$\Delta\boldsymbol{G} = \boldsymbol{G}\boldsymbol{P}^{-1}\Big(\underbrace{\mathrm{cov}(\boldsymbol{\mu})}_{\frac{\boldsymbol{P}}{2}\text{ (Eq. 10)}} - \frac{1}{2}\boldsymbol{P}\Big)\boldsymbol{P}^{-1}\boldsymbol{G} = 0. \tag{22}$$

The way to incorporate the effect of selection is again to assume that the mean midparent phenotype $\boldsymbol{\mu}$ has changed to $\tilde{\boldsymbol{\mu}}$. Using this and Eq. 10, this means we have to replace $\mathrm{cov}(\boldsymbol{\mu}) = \boldsymbol{P}/2$ with $\mathrm{cov}(\tilde{\boldsymbol{\mu}}) = \tilde{\boldsymbol{P}}/2$ in Eq. 22:

$$\Delta \boldsymbol{G} = \frac{1}{2}\boldsymbol{G}\boldsymbol{P}^{-1}(\tilde{\boldsymbol{P}} - \boldsymbol{P})\boldsymbol{P}^{-1}\boldsymbol{G}, \tag{23}$$

where $\tilde{\boldsymbol{P}}$ is the phenotypic covariance of parents after selection. Once again, the equation is now expressed through quantities pertaining to parents as opposed to midparents.

Note that this derivation of the genetic covariance under selection tacitly assumes that the change in segregation variance $\boldsymbol{S}$ from one generation to the next can be neglected (Taper and Case 1985). This is a good approximation however, echoed by many selection experiments.

#### 2 Community dynamics

We model the dynamics of $S$ species differing in $L$ ecologically relevant traits, collected in the vector $\boldsymbol{z}$. The trait distribution within each species $i$ is then

$$p_i(\boldsymbol{z}) = \mathcal{N}(\boldsymbol{z}; \boldsymbol{\mu}_i, \boldsymbol{P}_i) \tag{24}$$

(with $\mathcal{N}(\boldsymbol{z}; \boldsymbol{\mu}_i, \boldsymbol{P}_i)$ given by Eq. 1), where the mean trait $\boldsymbol{\mu}_i$ and phenotypic covariance matrix $\boldsymbol{P}_i$ are time-dependent. It satisfies the normalization condition

$$\int_{-\infty}^{+\infty}\int_{-\infty}^{+\infty}\cdots\int_{-\infty}^{+\infty} p_i(\boldsymbol{z})\,\mathrm{d}z^1\mathrm{d}z^2\cdots\mathrm{d}z^L = 1 \tag{25}$$

at every moment of time (superscripts show trait indices; subscripts are reserved for the species index). Using the notation $\mathrm{d}\boldsymbol{z} = \mathrm{d}z^1\mathrm{d}z^2\cdots\mathrm{d}z^L$, we write the integral simply as

$$\int p_i(\boldsymbol{z})\,\mathrm{d}\boldsymbol{z} = 1. \tag{26}$$

The total population density of species $i$ is $N_i$. Assuming discrete non-overlapping generations, the life cycle follows

$$N_i p_i(\boldsymbol{z}) \xrightarrow{\text{selection}} \tilde{N}_i \tilde{p}_i(\boldsymbol{z}) \xrightarrow{\text{reproduction}} N_i' p_i'(z), \tag{27}$$

where tildes denote state after selection but before reproduction, and primes the state in the next generation. Let $W_i(\boldsymbol{z})$ be the fitness of species $i$'s phenotype $\boldsymbol{z}$ at a given moment in time:

$$\tilde{N}_i \tilde{p}_i(\boldsymbol{z}) = W_i(\boldsymbol{z}) N_i p_i(\boldsymbol{z}). \tag{28}$$

$W_i(\boldsymbol{z})$ generally depends on the abundances and trait distributions of all interacting species. Here we apply the standard sleight of hand of assuming that $W_i(\boldsymbol{z})$ encodes both birth and death processes, and the only role of “reproduction” in Eq. 27 is to restore the trait distribution’s shape to normal—in other words, $\tilde{N}_i = N_i'$. The error made this way is negligible in the weak selection limit (Barabás and D’Andrea 2016). This limit means $W_i(\boldsymbol{z})$ can be written as

$$W_i(\boldsymbol{z}) = 1 + s r_i(\boldsymbol{z}), \tag{29}$$

where $s \ll 1$ is a small parameter and $r_i(\boldsymbol{z})$ is the per capita growth rate of species $i$, determined by the ecological interactions within the community.

The dynamics of the population densities from one generation to the next is obtained by integrating Eq. 28 over $\boldsymbol{z}$:

$$\tilde{N}_i \int \tilde{p}_i(\boldsymbol{z})\,\mathrm{d}\boldsymbol{z} = N_i \int W_i(\boldsymbol{z}) p_i(\boldsymbol{z})\,\mathrm{d}\boldsymbol{z}. \tag{30}$$

From Eq. 26, the integral on the left hand side is 1. Using $\tilde{N}_i = N_i'$,

$$N_i' = N_i \int W_i(\boldsymbol{z}) p_i(\boldsymbol{z})\,\mathrm{d}\boldsymbol{z}. \tag{31}$$

In the weak selection limit of Eq. 29,

$$N_i' = N_i \int (1 + s r_i(\boldsymbol{z})) p_i(\boldsymbol{z})\,\mathrm{d}\boldsymbol{z} = N_i \left( 1 + s \int r_i(\boldsymbol{z}) p_i(\boldsymbol{z})\,\mathrm{d}\boldsymbol{z} \right), \tag{32}$$

where Eq. 26 was used again in the last step. Subtracting $N_i$ and dividing by $s$ gives

$$\frac{N_i' - N_i}{s} = N_i \int r_i(\boldsymbol{z}) p_i(\boldsymbol{z})\,\mathrm{d}\boldsymbol{z}, \tag{33}$$

which, with appropriate scaling and for $s \to 0$, leads to the following differential equation (Barabás and D'Andrea 2016):

$$\frac{\mathrm{d}N_i}{\mathrm{d}t} = N_i \int r_i(\boldsymbol{z}) p_i(\boldsymbol{z})\,\mathrm{d}\boldsymbol{z}. \tag{34}$$

To track the change of trait means, we use the multidimensional breeder's equation, Eq. 21. To do so, we need to express $\tilde{\boldsymbol{\mu}}_i$. By definition, the mean of $p_i(\boldsymbol{z})$ is

$$\boldsymbol{\mu}_i = \int \boldsymbol{z} p_i(\boldsymbol{z})\,\mathrm{d}\boldsymbol{z}, \tag{35}$$

where the integral is interpreted entry-wise: $\mu_i^k = \int z^k p_i(\boldsymbol{z})\,\mathrm{d}\boldsymbol{z}$. Let us rearrange Eq. 28:

$$\tilde{p}_i(\boldsymbol{z}) = \frac{W_i(\boldsymbol{z}) N_i p_i(\boldsymbol{z})}{\tilde{N}_i}. \tag{36}$$

Using $\tilde{N}_i = N_i'$ and then Eq. 31 in the denominator:

$$\tilde{p}_i(\boldsymbol{z}) = \frac{W_i(\boldsymbol{z}) N_i p_i(\boldsymbol{z})}{N_i \int W_i(\boldsymbol{z}) p_i(\boldsymbol{z})\,\mathrm{d}\boldsymbol{z}} = \frac{W_i(\boldsymbol{z}) p_i(\boldsymbol{z})}{\int W_i(\boldsymbol{z}) p_i(\boldsymbol{z})\,\mathrm{d}\boldsymbol{z}}. \tag{37}$$

Multiplying both sides by $\boldsymbol{z}$ and integrating, we get

$$\int \boldsymbol{z} \tilde{p}_i(\boldsymbol{z})\,\mathrm{d}\boldsymbol{z} = \frac{\int \boldsymbol{z} W_i(\boldsymbol{z}) p_i(\boldsymbol{z})\,\mathrm{d}\boldsymbol{z}}{\int W_i(\boldsymbol{z}) p_i(\boldsymbol{z})\,\mathrm{d}\boldsymbol{z}}. \tag{38}$$

Applying Eq. 35 on the left, the integral is simply $\tilde{\boldsymbol{\mu}}_i$:

$$\tilde{\boldsymbol{\mu}}_i = \frac{\int \boldsymbol{z} W_i(\boldsymbol{z}) p_i(\boldsymbol{z})\,\mathrm{d}\boldsymbol{z}}{\int W_i(\boldsymbol{z}) p_i(\boldsymbol{z})\,\mathrm{d}\boldsymbol{z}}. \tag{39}$$

Substituting this into Eq. 21 gives

$$\Delta\boldsymbol{\mu}_i = \boldsymbol{G}_i\boldsymbol{P}_i^{-1}\left(\frac{\int \boldsymbol{z}W_i(\boldsymbol{z})p_i(\boldsymbol{z})\,\mathrm{d}\boldsymbol{z}}{\int W_i(\boldsymbol{z})p_i(\boldsymbol{z})\,\mathrm{d}\boldsymbol{z}} - \boldsymbol{\mu}_i\right). \tag{40}$$

We again apply the weak selection limit (Eq. 29):

$$\Delta\boldsymbol{\mu}_i = \boldsymbol{G}_i\boldsymbol{P}_i^{-1}\left(\frac{\int \boldsymbol{z}(1 + sr_i(\boldsymbol{z}))p_i(\boldsymbol{z})\,\mathrm{d}\boldsymbol{z}}{\int (1 + sr_i(\boldsymbol{z}))p_i(\boldsymbol{z})\,\mathrm{d}\boldsymbol{z}} - \boldsymbol{\mu}_i\right), \tag{41}$$

or

$$\Delta\boldsymbol{\mu}_i = \boldsymbol{G}_i\boldsymbol{P}_i^{-1}\left(\frac{\int \boldsymbol{z}p_i(\boldsymbol{z})\,\mathrm{d}\boldsymbol{z} + s\int \boldsymbol{z}r_i(\boldsymbol{z})p_i(\boldsymbol{z})\,\mathrm{d}\boldsymbol{z}}{1 + s\int r_i(\boldsymbol{z})p_i(\boldsymbol{z})\,\mathrm{d}\boldsymbol{z}} - \boldsymbol{\mu}_i\right). \tag{42}$$

The first integral in the numerator is $\boldsymbol{\mu}_i$ by Eq. 35:

$$\Delta\boldsymbol{\mu}_i = \boldsymbol{G}_i\boldsymbol{P}_i^{-1}\left(\frac{\boldsymbol{\mu}_i + s\int \boldsymbol{z}r_i(\boldsymbol{z})p_i(\boldsymbol{z})\,\mathrm{d}\boldsymbol{z}}{1 + s\int r_i(\boldsymbol{z})p_i(\boldsymbol{z})\,\mathrm{d}\boldsymbol{z}} - \boldsymbol{\mu}_i\right). \tag{43}$$

Taylor expanding to first order in $s$ gives

$$\Delta\boldsymbol{\mu}_i = \boldsymbol{G}_i\boldsymbol{P}_i^{-1}\left[\boldsymbol{\mu}_i + s\left(\int \boldsymbol{z}r_i(\boldsymbol{z})p_i(\boldsymbol{z})\,\mathrm{d}\boldsymbol{z} - \boldsymbol{\mu}_i\int r_i(\boldsymbol{z})p_i(\boldsymbol{z})\,\mathrm{d}\boldsymbol{z}\right) - \boldsymbol{\mu}_i\right] + O(s^2), \tag{44}$$

where $O(s^k)$ means terms proportional to the $k$th power of $s$ or higher. After simplifying,

$$\Delta\boldsymbol{\mu}_i = s\,\boldsymbol{G}_i\boldsymbol{P}_i^{-1}\int (\boldsymbol{z} - \boldsymbol{\mu}_i)r_i(\boldsymbol{z})p_i(\boldsymbol{z})\,\mathrm{d}\boldsymbol{z} + O(s^2). \tag{45}$$

Dividing by $s$ and performing the $s \to 0$ limit gives the differential equation

$$\frac{\mathrm{d}\boldsymbol{\mu}_i}{\mathrm{d}t} = \boldsymbol{G}_i\boldsymbol{P}_i^{-1}\int (\boldsymbol{z} - \boldsymbol{\mu}_i)r_i(\boldsymbol{z})p_i(\boldsymbol{z})\,\mathrm{d}\boldsymbol{z}. \tag{46}$$

Finally, the change in the genetic covariance matrices $\boldsymbol{G}_i$ can be tracked via Eq. 23. To use this equation, we first need to express $\tilde{\boldsymbol{P}}_i$. In general, the covariance matrix of a vector $\boldsymbol{x}$ reads

$$\mathrm{cov}(\boldsymbol{x}) = \int (\boldsymbol{z} - \boldsymbol{x}) \circ (\boldsymbol{z} - \boldsymbol{x})p_i(\boldsymbol{z})\,\mathrm{d}\boldsymbol{z}, \tag{47}$$

where $\circ$ denotes the outer product (i.e, for vectors $\boldsymbol{a}$ and $\boldsymbol{b}$ with entries $a^k$ and $b^k$, the matrix $\boldsymbol{A} = \boldsymbol{a} \circ \boldsymbol{b}$ has entries $A^{kl} = a^k b^l$). As before, the integral is interpreted entry-wise: $[\mathrm{cov}(\boldsymbol{x})]^{kl} = \int (z^k - x^k)(z^l - x^l)p_i(\boldsymbol{z})\,\mathrm{d}\boldsymbol{z}$. Based on this, $\tilde{\boldsymbol{P}}_i$ reads

$$\tilde{\boldsymbol{P}}_i = \int (\boldsymbol{z} - \tilde{\boldsymbol{\mu}}_i) \circ (\boldsymbol{z} - \tilde{\boldsymbol{\mu}}_i)\tilde{p}_i(\boldsymbol{z})\,\mathrm{d}\boldsymbol{z}, \tag{48}$$

or, using Eq. 37,

$$\tilde{\boldsymbol{P}}_i = \frac{\int (\boldsymbol{z} - \tilde{\boldsymbol{\mu}}_i) \circ (\boldsymbol{z} - \tilde{\boldsymbol{\mu}}_i)W_i(\boldsymbol{z})p_i(\boldsymbol{z})\,\mathrm{d}\boldsymbol{z}}{\int W_i(\boldsymbol{z})p_i(\boldsymbol{z})\,\mathrm{d}\boldsymbol{z}}. \tag{49}$$

In the weak selection limit (Eq. 29), this is written as

$$\tilde{\boldsymbol{P}}_i = \frac{\displaystyle\int (\boldsymbol{z} - \tilde{\boldsymbol{\mu}}_i) \circ (\boldsymbol{z} - \tilde{\boldsymbol{\mu}}_i)(1 + s r_i(\boldsymbol{z})) p_i(\boldsymbol{z}) \, \mathrm{d}\boldsymbol{z}}{\displaystyle\int (1 + s r_i(\boldsymbol{z})) p_i(\boldsymbol{z}) \, \mathrm{d}\boldsymbol{z}} = \frac{\displaystyle\int (\boldsymbol{z} - \tilde{\boldsymbol{\mu}}_i) \circ (\boldsymbol{z} - \tilde{\boldsymbol{\mu}}_i) p_i(\boldsymbol{z}) \, \mathrm{d}\boldsymbol{z} + s \int (\boldsymbol{z} - \tilde{\boldsymbol{\mu}}_i) \circ (\boldsymbol{z} - \tilde{\boldsymbol{\mu}}_i) r_i(\boldsymbol{z}) p_i(\boldsymbol{z}) \, \mathrm{d}\boldsymbol{z}}{\displaystyle 1 + s \int r_i(\boldsymbol{z}) p_i(\boldsymbol{z}) \, \mathrm{d}\boldsymbol{z}}. \tag{50}$$

Taylor expanding in $s$ to linear order yields

$$\tilde{\boldsymbol{P}}_i = \int (\boldsymbol{z} - \tilde{\boldsymbol{\mu}}_i) \circ (\boldsymbol{z} - \tilde{\boldsymbol{\mu}}_i) p_i(\boldsymbol{z}) \, \mathrm{d}\boldsymbol{z} + s \left[ \int (\boldsymbol{z} - \tilde{\boldsymbol{\mu}}_i) \circ (\boldsymbol{z} - \tilde{\boldsymbol{\mu}}_i) r_i(\boldsymbol{z}) p_i(\boldsymbol{z}) \, \mathrm{d}\boldsymbol{z} - \left( \int (\boldsymbol{z} - \tilde{\boldsymbol{\mu}}_i) \circ (\boldsymbol{z} - \tilde{\boldsymbol{\mu}}_i) p_i(\boldsymbol{z}) \, \mathrm{d}\boldsymbol{z} \right) \left( \int r_i(\boldsymbol{z}) p_i(\boldsymbol{z}) \, \mathrm{d}\boldsymbol{z} \right) \right] + O(s^2). \tag{51}$$

To make further progress, we need to express $\tilde{\boldsymbol{\mu}}_i$ via $\boldsymbol{\mu}_i$. This can be done by equating the right hand sides of Eqs. 21 and 45, and solving for $\tilde{\boldsymbol{\mu}}$, leading to

$$\tilde{\boldsymbol{\mu}}_i = \boldsymbol{\mu}_i + s \int (\boldsymbol{z} - \boldsymbol{\mu}_i) r_i(\boldsymbol{z}) p_i(\boldsymbol{z}) \, \mathrm{d}\boldsymbol{z} + O(s^2). \tag{52}$$

As a simplifying notation, let us call the integral above $\boldsymbol{q}_i$: $\tilde{\boldsymbol{\mu}}_i = \boldsymbol{\mu}_i + s\boldsymbol{q}_i + O(s^2)$. We substitute this into Eq. 51 and neglect higher-order terms in $s$. Let us first consider the first term in Eq. 51:

$$\int (\boldsymbol{z} - \tilde{\boldsymbol{\mu}}_i) \circ (\boldsymbol{z} - \tilde{\boldsymbol{\mu}}_i) p_i(\boldsymbol{z}) \, \mathrm{d}\boldsymbol{z} = \int (\boldsymbol{z} - \boldsymbol{\mu}_i - s\boldsymbol{q}_i) \circ (\boldsymbol{z} - \boldsymbol{\mu}_i - s\boldsymbol{q}_i) p_i(\boldsymbol{z}) \, \mathrm{d}\boldsymbol{z} + O(s^2) = \underbrace{\int (\boldsymbol{z} - \boldsymbol{\mu}_i) \circ (\boldsymbol{z} - \boldsymbol{\mu}_i) p_i(\boldsymbol{z}) \, \mathrm{d}\boldsymbol{z}}_{\boldsymbol{P}_i} - 2s \underbrace{\left( \int (\boldsymbol{z} - \boldsymbol{\mu}_i) p_i(\boldsymbol{z}) \, \mathrm{d}\boldsymbol{z} \right)}_{0} \circ \boldsymbol{q}_i + O(s^2) = \boldsymbol{P}_i + O(s^2). \tag{53}$$

The first braced term is by definition the total phenotypic covariance, while the second is zero because it is the integral of an odd and an even function in $\boldsymbol{z} - \boldsymbol{\mu}_i$ (Eq. 24). The second term in Eq. 51 is much simpler: since this is already proportional to $s$, any $s\boldsymbol{q}_i$ terms will be proportional to $s^2$ and can be neglected. We can therefore simply omit the $s\boldsymbol{q}_i$ terms, which means replacing $\tilde{\boldsymbol{\mu}}_i$ by $\boldsymbol{\mu}_i$. The second term in Eq. 51 therefore reads

$$s \left[ \int (\boldsymbol{z} - \tilde{\boldsymbol{\mu}}_i) \circ (\boldsymbol{z} - \tilde{\boldsymbol{\mu}}_i) r_i(\boldsymbol{z}) p_i(\boldsymbol{z}) \, \mathrm{d}\boldsymbol{z} - \left( \int (\boldsymbol{z} - \tilde{\boldsymbol{\mu}}_i) \circ (\boldsymbol{z} - \tilde{\boldsymbol{\mu}}_i) p_i(\boldsymbol{z}) \, \mathrm{d}\boldsymbol{z} \right) \left( \int r_i(\boldsymbol{z}) p_i(\boldsymbol{z}) \, \mathrm{d}\boldsymbol{z} \right) \right] = s \left[ \int (\boldsymbol{z} - \boldsymbol{\mu}_i) \circ (\boldsymbol{z} - \boldsymbol{\mu}_i) r_i(\boldsymbol{z}) p_i(\boldsymbol{z}) \, \mathrm{d}\boldsymbol{z} - \underbrace{\left( \int (\boldsymbol{z} - \boldsymbol{\mu}_i) \circ (\boldsymbol{z} - \boldsymbol{\mu}_i) p_i(\boldsymbol{z}) \, \mathrm{d}\boldsymbol{z} \right)}_{\boldsymbol{P}_i} \left( \int r_i(\boldsymbol{z}) p_i(\boldsymbol{z}) \, \mathrm{d}\boldsymbol{z} \right) \right] + O(s^2), \tag{54}$$

or

$$s \left[ \int (\boldsymbol{z} - \boldsymbol{\mu}_i) \circ (\boldsymbol{z} - \boldsymbol{\mu}_i) r_i(\boldsymbol{z}) p_i(\boldsymbol{z}) \, \mathrm{d}\boldsymbol{z} - \boldsymbol{P}_i \int r_i(\boldsymbol{z}) p_i(\boldsymbol{z}) \, \mathrm{d}\boldsymbol{z} \right] + O(s^2) = s \int [(\boldsymbol{z} - \boldsymbol{\mu}_i) \circ (\boldsymbol{z} - \boldsymbol{\mu}_i) - \boldsymbol{P}_i] r_i(\boldsymbol{z}) p_i(\boldsymbol{z}) \, \mathrm{d}\boldsymbol{z} + O(s^2). \tag{55}$$

Substituting Eqs. 53 and 54 into Eq. 51:

$$\tilde{\boldsymbol{P}}_i = \boldsymbol{P}_i + s \int [(\boldsymbol{z} - \boldsymbol{\mu}_i) \circ (\boldsymbol{z} - \boldsymbol{\mu}_i) - \boldsymbol{P}_i] r_i(\boldsymbol{z}) p_i(\boldsymbol{z}) \,\mathrm{d}\boldsymbol{z} + O(s^2). \tag{56}$$

Substituting this in turn into Eq. 23, we get

$$\begin{aligned} \Delta \boldsymbol{G}_i &= \frac{1}{2} \boldsymbol{G}_i \boldsymbol{P}_i^{-1} \left[ \boldsymbol{P}_i + s \int [(\boldsymbol{z} - \boldsymbol{\mu}_i) \circ (\boldsymbol{z} - \boldsymbol{\mu}_i) - \boldsymbol{P}_i] r_i(\boldsymbol{z}) p_i(\boldsymbol{z}) \,\mathrm{d}\boldsymbol{z} - \boldsymbol{P}_i \right] \boldsymbol{P}_i^{-1} \boldsymbol{G}_i + O(s^2) \\ &= s \frac{1}{2} \boldsymbol{G}_i \boldsymbol{P}_i^{-1} \left[ \int [(\boldsymbol{z} - \boldsymbol{\mu}_i) \circ (\boldsymbol{z} - \boldsymbol{\mu}_i) - \boldsymbol{P}_i] r_i(\boldsymbol{z}) p_i(\boldsymbol{z}) \,\mathrm{d}\boldsymbol{z} \right] \boldsymbol{P}_i^{-1} \boldsymbol{G}_i + O(s^2). \end{aligned} \tag{57}$$

Dividing by $s$ and taking $s \to 0$ leads to the differential equation

$$\frac{\mathrm{d}\boldsymbol{G}_i}{\mathrm{d}t} = \frac{1}{2} \boldsymbol{G}_i \boldsymbol{P}_i^{-1} \left[ \int [(\boldsymbol{z} - \boldsymbol{\mu}_i) \circ (\boldsymbol{z} - \boldsymbol{\mu}_i) - \boldsymbol{P}_i] r_i(\boldsymbol{z}) p_i(\boldsymbol{z}) \,\mathrm{d}\boldsymbol{z} \right] \boldsymbol{P}_i^{-1} \boldsymbol{G}_i. \tag{58}$$

This concludes the derivation for the governing equations of the population densities, trait means, and trait covariances of the species, given by Eqs. 34, 46, and 58. Writing them out all together:

$$\frac{\mathrm{d}N_i}{\mathrm{d}t} = N_i \int r_i(\boldsymbol{z}) p_i(\boldsymbol{z}) \,\mathrm{d}\boldsymbol{z}, \tag{59}$$

$$\frac{\mathrm{d}\boldsymbol{\mu}_i}{\mathrm{d}t} = \boldsymbol{G}_i \boldsymbol{P}_i^{-1} \int (\boldsymbol{z} - \boldsymbol{\mu}_i) r_i(\boldsymbol{z}) p_i(\boldsymbol{z}) \,\mathrm{d}\boldsymbol{z}, \tag{60}$$

$$\frac{\mathrm{d}\boldsymbol{G}_i}{\mathrm{d}t} = \frac{1}{2} \boldsymbol{G}_i \boldsymbol{P}_i^{-1} \left[ \int [(\boldsymbol{z} - \boldsymbol{\mu}_i) \circ (\boldsymbol{z} - \boldsymbol{\mu}_i) - \boldsymbol{P}_i] r_i(\boldsymbol{z}) p_i(\boldsymbol{z}) \,\mathrm{d}\boldsymbol{z} \right] \boldsymbol{P}_i^{-1} \boldsymbol{G}_i. \tag{61}$$

### 3 The consumer-resource model

Eqs. 59, 60, and 61 only specify the dynamics if the per capita growth rates are given. Here we obtain these from a simple consumer-resource system. We have a set of resources which can be mapped onto an $L$-dimensional trait space. For instance, if $L = 1$ and the trait is the bill depth of Darwin's finches, then the resources are of various sizes; the resource which a finch of bill depth $\boldsymbol{y}$ can optimally utilize is what we call a resource of size $\boldsymbol{y}$. Let $R(\boldsymbol{y})$ be the availability of resource $\boldsymbol{y}$. The degree to which an individual of phenotype $\boldsymbol{z}$ can utilize resource $\boldsymbol{y}$ is given by $u(\boldsymbol{z}, \boldsymbol{y})$. The amount of resource available is the maximum (saturation) concentration of the resource, $R_0(\boldsymbol{y})$, minus what has been consumed:

$$R(\boldsymbol{y}) = R_0(\boldsymbol{y}) - \sum_{j=1}^{S} \int u(\boldsymbol{z}', \boldsymbol{y}) N_j p_j(\boldsymbol{z}') \,\mathrm{d}\boldsymbol{z}', \tag{62}$$

where $S$ is the number of consumer species, and $N_j p_j(\boldsymbol{z}')$ is the population density of consumer $j$'s individuals that have phenotype $\boldsymbol{z}'$. In turn, the per capita growth rate $r(\boldsymbol{z})$ of consumers with phenotype $\boldsymbol{z}$ is proportional to their total resource consumption, and to a phenotype-specific mortality rate $m(\boldsymbol{z})$:

$$r(\boldsymbol{z}) = \int u(\boldsymbol{z}, \boldsymbol{y}) R(\boldsymbol{y}) \,\mathrm{d}y - m(\boldsymbol{z}). \tag{63}$$

The fact that $R(\boldsymbol{y})$ is directly expressed means the resources are assumed to operate on a fast time scale compared to the population dynamics (MacArthur 1970), and are therefore always in a state of quasi-equilibrium. Also, resource depletion is weighted by the same function, $u(\boldsymbol{z}, \boldsymbol{y})$, as population growth in Eq. 63. this means that the benefit an individual gains from resource $\boldsymbol{y}$ is proportional to its consumption of the same resource.

Substituting Eq. 62 into Eq. 63 yields

$$r(\boldsymbol{z}) = \int u(\boldsymbol{z},\boldsymbol{y})\left(R_0(\boldsymbol{y}) - \sum_{j=1}^{S} \int u(\boldsymbol{z}',\boldsymbol{y}) N_j p_j(\boldsymbol{z}')\,\mathrm{d}\boldsymbol{z}'\right)\mathrm{d}\boldsymbol{y} - m(\boldsymbol{z}). \tag{64}$$

Rearranging, we get

$$r(\boldsymbol{z}) = \underbrace{\left(\int u(\boldsymbol{z},\boldsymbol{y})R_0(\boldsymbol{y})\,\mathrm{d}\boldsymbol{y} - m(\boldsymbol{z})\right)}_{b(\boldsymbol{z})} - \sum_{j=1}^{S} \int \underbrace{\left(\int u(\boldsymbol{z},\boldsymbol{y})u(\boldsymbol{z}',\boldsymbol{y})\,\mathrm{d}\boldsymbol{y}\right)}_{a(\boldsymbol{z},\boldsymbol{z}')} N_j p_j(\boldsymbol{z}')\,\mathrm{d}\boldsymbol{z}'. \tag{65}$$

With the definitions

$$b(\boldsymbol{z}) = \int u(\boldsymbol{z},\boldsymbol{y})R_0(\boldsymbol{y})\,\mathrm{d}\boldsymbol{y} - m(\boldsymbol{z}), \tag{66}$$

$$a(\boldsymbol{z},\boldsymbol{z}') = \int u(\boldsymbol{z},\boldsymbol{y})u(\boldsymbol{z}',\boldsymbol{y})\,\mathrm{d}\boldsymbol{y}, \tag{67}$$

this has the form of Lotka–Volterra growth with intrinsic rates $b(\boldsymbol{z})$ and competition kernel $a(\boldsymbol{z},\boldsymbol{z}')$:

$$r(\boldsymbol{z}) = b(\boldsymbol{z}) - \sum_{j=1}^{S} N_j \int a(\boldsymbol{z},\boldsymbol{z}')p_j(\boldsymbol{z}')\,\mathrm{d}\boldsymbol{z}'. \tag{68}$$

Assigning parameters to this model, the resource utilization curve $u(\boldsymbol{z},\boldsymbol{y})$ is a Gaussian function of the difference between consumer phenotype $\boldsymbol{z}$ and resource quality $\boldsymbol{y}$. We write it as

$$u(\boldsymbol{z},\boldsymbol{y}) = \left[(2\pi)^L \det(2\boldsymbol{W})\right]^{1/4} \mathcal{N}(\boldsymbol{z};\boldsymbol{y},\boldsymbol{W}), \tag{69}$$

where $\mathcal{N}$ is given by Eq. 1, and the factor in front is introduced for convenience, to simplify the expression for the competition kernel $a(\boldsymbol{z},\boldsymbol{z}')$ between phenotypes $\boldsymbol{z}$ and $\boldsymbol{z}'$ in the future. The matrix $\boldsymbol{W}$ is the effective width of the utilization curve (the within-phenotype niche width of Taper and Case 1985), determining the range of resources that an individual with phenotype $\boldsymbol{z}$ can access.

The competition kernel $a(\boldsymbol{z},\boldsymbol{z}')$ now reads, from Eq. 67:

$$a(\boldsymbol{z},\boldsymbol{z}') = \int u(\boldsymbol{z},\boldsymbol{y})u(\boldsymbol{z}',\boldsymbol{y})\,\mathrm{d}\boldsymbol{y} = \left[(2\pi)^L \det(2\boldsymbol{W})\right]^{1/2} \int \mathcal{N}(\boldsymbol{z}-\boldsymbol{y};\boldsymbol{0},\boldsymbol{W})\mathcal{N}(\boldsymbol{y};\boldsymbol{z}',\boldsymbol{W})\,\mathrm{d}\boldsymbol{y}. \tag{70}$$

The integral is the convolution of two normal functions, which is also normal with the summed means and covariances. We therefore have

$$a(\boldsymbol{z},\boldsymbol{z}') = \sqrt{(2\pi)^L \det(2\boldsymbol{W})}\,\mathcal{N}(\boldsymbol{z};\boldsymbol{z}',2\boldsymbol{W}). \tag{71}$$

Introducing the notation $\boldsymbol{\Omega} = 2\boldsymbol{W}$ and using Eq. 1, this simplifies to

$$a(\boldsymbol{z},\boldsymbol{z}') = \exp\left(-\frac{1}{2}(\boldsymbol{z}-\boldsymbol{y})\boldsymbol{\Omega}^{-1}(\boldsymbol{z}-\boldsymbol{y})\right). \tag{72}$$

To determine $b(\boldsymbol{z})$ (Eq. 66), we need to know the resource saturation densities $R_0(\boldsymbol{y})$ and the mortality rates $m(\boldsymbol{z})$. Generally, we are not concerned about their exact form, as long as they bound the region of growth for the species—either because $R_0(\boldsymbol{y})$ drops to zero at some point, or $m(\boldsymbol{z})$ becomes prohibitively large (or both). One possible parameterization proceeds by assuming that each resource saturates at the same level without consumption, and mortality increases with distance from a certain optimal point, which we take to be at $\boldsymbol{z} = \boldsymbol{0}$ without loss of generality. We thus define $R_0(\boldsymbol{y})$ as a constant, whose form is chosen for convenience to be

$$R_0(\boldsymbol{y}) = \text{const.} = \left[(2\pi)^L \det(\boldsymbol{\Omega})\right]^{-1/4}. \tag{73}$$

This way, the first term in $b(\boldsymbol{z})$ (Eq. 66) is simply equal to 1:

$$\int u(\boldsymbol{z}, \boldsymbol{y}) R_0(\boldsymbol{y})\, \mathrm{d}\boldsymbol{y} = \left[(2\pi)^L \det(\boldsymbol{\Omega})\right]^{-1/4} \int u(\boldsymbol{z}, \boldsymbol{y})\, \mathrm{d}\boldsymbol{y}, \tag{74}$$

which, by Eq. 69 and the fact that the normal distribution integrates to unity, is

$$\left[(2\pi)^L \det(\boldsymbol{\Omega})\right]^{-1/4} \int u(\boldsymbol{z}, \boldsymbol{y})\, \mathrm{d}\boldsymbol{y} = \int \mathcal{N}(\boldsymbol{z}; \boldsymbol{y}, \boldsymbol{\Omega}/2)\, \mathrm{d}\boldsymbol{y} = 1. \tag{75}$$

The mortality $m(\boldsymbol{z})$ increases with increasing distance from $\boldsymbol{z} = \boldsymbol{0}$. We will use two possible forms: either a quadratically increasing mortality with $m(\boldsymbol{z}) = (\boldsymbol{z}\boldsymbol{z})/\theta^2$, or the quartic function $(\boldsymbol{z}) = (\boldsymbol{z}\boldsymbol{z})^4/\theta^4$. Here $(\boldsymbol{z}\boldsymbol{z})$ is the scalar product of $\boldsymbol{z}$ with itself, equal to the squared length of $\boldsymbol{z}$. Using Eq. 75 and these expressions for the mortalities, we now obtain the intrinsic rates $b(\boldsymbol{z})$ from its definition in Eq. 66:

$$b(z) = \int u(\boldsymbol{z}, \boldsymbol{y}) R_0(\boldsymbol{y})\, \mathrm{d}\boldsymbol{y} - m(\boldsymbol{z}) = 1 - m(\boldsymbol{z}) = \begin{cases} 1 - \dfrac{(\boldsymbol{z}\boldsymbol{z})}{\theta^2} & \text{(quadratic mortality)} \\ 1 - \dfrac{(\boldsymbol{z}\boldsymbol{z})^2}{\theta^4} & \text{(quartic mortality)} \end{cases} \tag{76}$$

Without loss of generality, we fix $\theta = 1/2$. With this choice, positive growth can only be achieved in a sphere of diameter 1 in trait space. This means that all quantities with units of trait distance are measured relative to this diameter.

### 4 Eco-evolutionary dynamics in the consumer-resource model

We start from the per capita growth rates of Eq. 68. Substituting this into our general eco-evolutionary dynamics (Eqs. 59-61), we get

$$\frac{\mathrm{d}N_i}{\mathrm{d}t} = N_i \left[ \int b(\boldsymbol{z}) p_i(\boldsymbol{z})\, \mathrm{d}\boldsymbol{z} - \sum_{j=1}^{S} N_j \iint p_i(\boldsymbol{z}) a(\boldsymbol{z}, \boldsymbol{z}') p_j(\boldsymbol{z}')\, \mathrm{d}\boldsymbol{z}'\, \mathrm{d}\boldsymbol{z} \right], \tag{77}$$

$$\frac{\mathrm{d}\boldsymbol{\mu}_i}{\mathrm{d}t} = \boldsymbol{G}_i \boldsymbol{P}_i^{-1} \left[ \int (\boldsymbol{z} - \boldsymbol{\mu}_i) b(\boldsymbol{z}) p_i(\boldsymbol{z})\, \mathrm{d}\boldsymbol{z} - \sum_{j=1}^{S} N_j \iint (\boldsymbol{z} - \boldsymbol{\mu}_i) p_i(\boldsymbol{z}) a(\boldsymbol{z}, \boldsymbol{z}') p_j(\boldsymbol{z}')\, \mathrm{d}\boldsymbol{z}'\, \mathrm{d}\boldsymbol{z} \right], \tag{78}$$

$$\frac{\mathrm{d}\boldsymbol{G}_i}{\mathrm{d}t} = \frac{1}{2} \boldsymbol{G}_i \boldsymbol{P}_i^{-1} \Bigg[ \int [(\boldsymbol{z} - \boldsymbol{\mu}_i) \circ (\boldsymbol{z} - \boldsymbol{\mu}_i) - \boldsymbol{P}_i] b(\boldsymbol{z}) p_i(\boldsymbol{z})\, \mathrm{d}\boldsymbol{z} \tag{79}$$
$$- \sum_{j=1}^{S} N_j \iint [(\boldsymbol{z} - \boldsymbol{\mu}_i) \circ (\boldsymbol{z} - \boldsymbol{\mu}_i) - \boldsymbol{P}_i] p_i(\boldsymbol{z}) a(\boldsymbol{z}, \boldsymbol{z}') p_j(\boldsymbol{z}')\, \mathrm{d}\boldsymbol{z}'\, \mathrm{d}\boldsymbol{z} \Bigg] \boldsymbol{P}_i^{-1} \boldsymbol{G}_i.$$

We introduce some simplifying notation:

$$b_i = \int b(\boldsymbol{z}) p_i(\boldsymbol{z})\, \mathrm{d}\boldsymbol{z}, \tag{80}$$

$$\alpha_{ij} = \iint p_i(\boldsymbol{z}) a(\boldsymbol{z}, \boldsymbol{z}') p_j(\boldsymbol{z}') \, \mathrm{d}\boldsymbol{z}' \, \mathrm{d}\boldsymbol{z}, \tag{81}$$

$$\boldsymbol{g}_i = \int \boldsymbol{P}_i^{-1}(\boldsymbol{z} - \boldsymbol{\mu}_i) b(\boldsymbol{z}) p_i(\boldsymbol{z}) \, \mathrm{d}\boldsymbol{z}, \tag{82}$$

$$\boldsymbol{\beta}_{ij} = \iint \boldsymbol{P}_i^{-1}(\boldsymbol{z} - \boldsymbol{\mu}_i) p_i(\boldsymbol{z}) a(\boldsymbol{z}, \boldsymbol{z}') p_j(\boldsymbol{z}') \, \mathrm{d}\boldsymbol{z}' \, \mathrm{d}\boldsymbol{z}, \tag{83}$$

$$\boldsymbol{Q}_i = \int \boldsymbol{P}_i^{-1}[(\boldsymbol{z} - \boldsymbol{\mu}_i) \circ (\boldsymbol{z} - \boldsymbol{\mu}_i) - \boldsymbol{P}_i] \boldsymbol{P}_i^{-1} b(\boldsymbol{z}) p_i(\boldsymbol{z}) \, \mathrm{d}\boldsymbol{z}, \tag{84}$$

$$\boldsymbol{\Gamma}_{ij} = \iint \boldsymbol{P}_i^{-1}[(\boldsymbol{z} - \boldsymbol{\mu}_i) \circ (\boldsymbol{z} - \boldsymbol{\mu}_i) - \boldsymbol{P}_i] \boldsymbol{P}_i^{-1} p_i(\boldsymbol{z}) a(\boldsymbol{z}, \boldsymbol{z}') p_j(\boldsymbol{z}') \, \mathrm{d}\boldsymbol{z}' \, \mathrm{d}\boldsymbol{z}, \tag{85}$$

where $\boldsymbol{g}_i$ and $\boldsymbol{\beta}_{ij}$ are vectors, while $\boldsymbol{Q}_i$ and $\boldsymbol{\Gamma}_{ij}$ are matrices in trait space. With these notations, Eqs. 77-79 are written as

$$\frac{\mathrm{d}N_i}{\mathrm{d}t} = N_i \left( b_i - \sum_{j=1}^{S} \alpha_{ij} N_j \right), \tag{86}$$

$$\frac{\mathrm{d}\boldsymbol{\mu}_i}{\mathrm{d}t} = \boldsymbol{G}_i \left( \boldsymbol{g}_i - \sum_{j=1}^{S} \boldsymbol{\beta}_{ij} N_j \right), \tag{87}$$

$$\frac{\mathrm{d}\boldsymbol{G}_i}{\mathrm{d}t} = \frac{1}{2} \boldsymbol{G}_i \left( \boldsymbol{Q}_i - \sum_{j=1}^{S} \boldsymbol{\Gamma}_{ij} N_j \right) \boldsymbol{G}_i. \tag{88}$$

To further relate Eqs. 80-85 to one another, we calculate the first two derivatives of the normal distribution with respect to its mean. From Eq. 1, the first derivative reads

$$\begin{aligned}
\frac{\partial \mathcal{N}(\boldsymbol{z}; \boldsymbol{\mu}, \boldsymbol{P})}{\partial \boldsymbol{\mu}} &= \frac{\partial \mathcal{N}(\boldsymbol{z}; \boldsymbol{\mu}, \boldsymbol{P})}{\partial \mu^k} = -\frac{1}{2} \mathcal{N}(\boldsymbol{z}; \boldsymbol{\mu}, \boldsymbol{P}) \frac{\partial}{\partial \mu^k} \sum_{m,n} (P^{-1})^{mn} (z^m - \mu^m)(z^n - \mu^n) \\
&= \frac{1}{2} \mathcal{N}(\boldsymbol{z}; \boldsymbol{\mu}, \boldsymbol{P}) \sum_{m,n} (P^{-1})^{mn} (z^m \delta^{nk} + z^n \delta^{mk} - \delta^{mk} \mu^n - \delta^{nk} \mu^m) \\
&= \frac{1}{2} \mathcal{N}(\boldsymbol{z}; \boldsymbol{\mu}, \boldsymbol{P}) \sum_{m,n} \left[ (P^{-1})^{mk} z^m + (P^{-1})^{kn} z^n - (P^{-1})^{kn} \mu^n - (P^{-1})^{mk} \mu^m \right] \\
&= \mathcal{N}(\boldsymbol{z}; \boldsymbol{\mu}, \boldsymbol{P}) \sum_{m} (P^{-1})^{km} (z^m - \mu^m) \\
&= \mathcal{N}(\boldsymbol{z}; \boldsymbol{\mu}, \boldsymbol{P}) \boldsymbol{P}^{-1} (\boldsymbol{z} - \boldsymbol{\mu}),
\end{aligned} \tag{89}$$

where $\mu^k$ is the $k$th trait-component of the vector $\boldsymbol{\mu}$, $(P^{-1})^{mn}$ is the $(m, n)$th entry of the matrix $\boldsymbol{P}^{-1}$, summations go from 1 to the number of trait dimensions $L$, and $\delta^{kl}$ is the Kronecker symbol (equal to 1 if $k = l$ and to 0 otherwise). Since $\mathcal{N}(\boldsymbol{z}; \boldsymbol{\mu}, \boldsymbol{P}) = \mathcal{N}(\boldsymbol{\mu}; \boldsymbol{z}, \boldsymbol{P})$ (Eq. 1), the derivative with respect to the independent variable (as opposed to the mean) immediately follows from Eq. 89:

$$\frac{\partial \mathcal{N}(\boldsymbol{z}; \boldsymbol{\mu}, \boldsymbol{P})}{\partial \boldsymbol{z}} = \mathcal{N}(\boldsymbol{z}; \boldsymbol{\mu}, \boldsymbol{P}) \boldsymbol{P}^{-1} (\boldsymbol{\mu} - \boldsymbol{z}). \tag{90}$$

Similarly, the second derivative with respect to the mean reads

$$
\begin{aligned}
\frac{\partial^2 \mathcal{N}(\boldsymbol{z};\boldsymbol{\mu},\boldsymbol{P})}{\partial\boldsymbol{\mu}\circ\partial\boldsymbol{\mu}} &= \frac{\partial^2 \mathcal{N}(\boldsymbol{z};\boldsymbol{\mu},\boldsymbol{P})}{\partial\mu^k\partial\mu^l} = \frac{\partial}{\partial\mu^k}\left[\frac{\partial \mathcal{N}(\boldsymbol{z};\boldsymbol{\mu},\boldsymbol{P})}{\partial\mu^l}\right] \\
&= \frac{\partial}{\partial\mu^k}\left[\mathcal{N}(\boldsymbol{z};\boldsymbol{\mu},\boldsymbol{P})\sum_m (P^{-1})^{lm}(z^m-\mu^m)\right] \\
&= \frac{\partial \mathcal{N}(\boldsymbol{z};\boldsymbol{\mu},\boldsymbol{P})}{\partial\mu^k}\sum_m (P^{-1})^{lm}(z^m-\mu^m) + \mathcal{N}(\boldsymbol{z};\boldsymbol{\mu},\boldsymbol{P})\left[-\sum_m (P^{-1})^{lm}\delta^{mk}\right] \\
&= \mathcal{N}(\boldsymbol{z};\boldsymbol{\mu},\boldsymbol{P})\left[\sum_m (P^{-1})^{km}(z^m-\mu^m)\right]\left[\sum_m (P^{-1})^{lm}(z^m-\mu^m)\right] - \mathcal{N}(\boldsymbol{z};\boldsymbol{\mu},\boldsymbol{P})(P^{-1})^{lk} \\
&= \mathcal{N}(\boldsymbol{z};\boldsymbol{\mu},\boldsymbol{P})\left[\left(\boldsymbol{P}^{-1}(\boldsymbol{z}-\boldsymbol{\mu})\right)\circ\left(\boldsymbol{P}^{-1}(\boldsymbol{z}-\boldsymbol{\mu})\right) - \boldsymbol{P}^{-1}\right] \\
&= \mathcal{N}(\boldsymbol{z};\boldsymbol{\mu},\boldsymbol{P})\boldsymbol{P}^{-1}[(\boldsymbol{z}-\boldsymbol{\mu})\circ(\boldsymbol{z}-\boldsymbol{\mu}) - \boldsymbol{P}]\boldsymbol{P}^{-1},
\end{aligned} \tag{91}
$$

where we used the fact that $\boldsymbol{P}$, and therefore its inverse, is symmetric. From $\mathcal{N}(\boldsymbol{z};\boldsymbol{\mu},\boldsymbol{P}) = \mathcal{N}(\boldsymbol{\mu};\boldsymbol{z},\boldsymbol{P})$ again, the second derivative with respect to the independent variable is the same,

$$
\frac{\partial^2 \mathcal{N}(\boldsymbol{z};\boldsymbol{\mu},\boldsymbol{P})}{\partial\boldsymbol{z}\circ\partial\boldsymbol{z}} = \mathcal{N}(\boldsymbol{z};\boldsymbol{\mu},\boldsymbol{P})\boldsymbol{P}^{-1}[(\boldsymbol{z}-\boldsymbol{\mu})\circ(\boldsymbol{z}-\boldsymbol{\mu}) - \boldsymbol{P}]\boldsymbol{P}^{-1}. \tag{92}
$$

With these results and the definitions in Eqs. 80-85, we can relate $\boldsymbol{g}_i$, $\boldsymbol{\beta}_{ij}$, $\boldsymbol{Q}_i$, and $\boldsymbol{\Gamma}_{ij}$ to derivatives of $b_i$ and $\alpha_{ij}$, keeping in mind that $p_i(\boldsymbol{z})$ is given by Eq. 24:

$$
\frac{\partial b_i}{\partial\boldsymbol{\mu}_i} = \frac{\partial}{\partial\boldsymbol{\mu}_i}\int b(\boldsymbol{z})p_i(\boldsymbol{z})\,\mathrm{d}\boldsymbol{z} = \int b(\boldsymbol{z})\frac{\partial p_i(\boldsymbol{z})}{\partial\boldsymbol{\mu}_i}\,\mathrm{d}\boldsymbol{z} = \int \boldsymbol{P}_i^{-1}(\boldsymbol{z}-\boldsymbol{\mu}_i)b(\boldsymbol{z})p_i(\boldsymbol{z}) = \boldsymbol{g}_i, \tag{93}
$$

$$
\frac{\partial\alpha_{ij}}{\partial\boldsymbol{\mu}_i} = \iint \frac{\partial p_i(\boldsymbol{z})}{\partial\boldsymbol{\mu}_i}a(\boldsymbol{z},\boldsymbol{z}')p_j(\boldsymbol{z}')\,\mathrm{d}\boldsymbol{z}'\,\mathrm{d}\boldsymbol{z} = \iint \boldsymbol{P}_i^{-1}(\boldsymbol{z}-\boldsymbol{\mu}_i)p_i(\boldsymbol{z})a(\boldsymbol{z},\boldsymbol{z}')p_j(\boldsymbol{z}')\,\mathrm{d}\boldsymbol{z}'\,\mathrm{d}\boldsymbol{z} = \boldsymbol{\beta}_{ij}, \tag{94}
$$

$$
\begin{aligned}
\frac{\partial^2 b_i}{\partial\boldsymbol{\mu}_i\circ\partial\boldsymbol{\mu}_i} &= \frac{1}{2}\int b(\boldsymbol{z})\frac{\partial^2 p_i(\boldsymbol{z})}{\partial\boldsymbol{\mu}_i\circ\partial\boldsymbol{\mu}_i}\,\mathrm{d}\boldsymbol{z} \\
&= \frac{1}{2}\int b(\boldsymbol{z})p_i(\boldsymbol{z})\boldsymbol{P}_i^{-1}[(\boldsymbol{z}-\boldsymbol{\mu}_i)\circ(\boldsymbol{z}-\boldsymbol{\mu}_i) - \boldsymbol{P}_i]\boldsymbol{P}_i^{-1}\,\mathrm{d}\boldsymbol{z} = \boldsymbol{Q}_i,
\end{aligned} \tag{95}
$$

$$
\begin{aligned}
\frac{\partial^2\alpha_{ij}}{\partial\boldsymbol{\mu}_i\circ\partial\boldsymbol{\mu}_i} &= \iint \frac{\partial p_i(\boldsymbol{z})}{\partial\boldsymbol{\mu}_i\circ\partial\boldsymbol{\mu}_i}a(\boldsymbol{z},\boldsymbol{z}')p_j(\boldsymbol{z}')\,\mathrm{d}\boldsymbol{z}\,\mathrm{d}\boldsymbol{z}' \\
&= \iint \boldsymbol{P}_i^{-1}[(\boldsymbol{z}-\boldsymbol{\mu}_i)\circ(\boldsymbol{z}-\boldsymbol{\mu}_i) - \boldsymbol{P}_i]\boldsymbol{P}_i^{-1}p_i(\boldsymbol{z})a(\boldsymbol{z},\boldsymbol{z}')p_i(\boldsymbol{z}')\,\mathrm{d}\boldsymbol{z}'\,\mathrm{d}\boldsymbol{z} = \boldsymbol{\Gamma}_{ij}.
\end{aligned} \tag{96}
$$

Note that $\partial\alpha_{ij}/\partial\boldsymbol{\mu}_i$ is interpreted as $\partial_1\alpha(\boldsymbol{\mu}_i,\boldsymbol{\mu}_j)$, the derivative with respect to the first argument. This means that even when $\boldsymbol{\mu}_j = \boldsymbol{\mu}_i$, the derivative is taken only with respect to the first $\boldsymbol{\mu}_i$, treating $\boldsymbol{\mu}_j = \boldsymbol{\mu}_i$ formally distinct.

We can now determine the functions in Eqs. 80-85 explicitly, based on the forms of $a(\boldsymbol{z},\boldsymbol{z}')$ (Eq. 72) and $b(\boldsymbol{z})$ (Eq. 76). We have, from the definition of $\alpha_{ij}$ in Eq. 81,

$$
\alpha_{ij} = \iint p_i(\boldsymbol{z})a(\boldsymbol{z},\boldsymbol{z}')p_j(\boldsymbol{z}')\,\mathrm{d}\boldsymbol{z}'\,\mathrm{d}\boldsymbol{z} = \int p_i(\boldsymbol{z})\left(\int a(\boldsymbol{z},\boldsymbol{z}')p_j(\boldsymbol{z}')\,\mathrm{d}\boldsymbol{z}'\right)\mathrm{d}\boldsymbol{z}. \tag{97}
$$

The inner integral can be written as the convolution of two multinormal distributions:

$$
\begin{aligned}
\int a(\boldsymbol{z},\boldsymbol{z}')p_j(\boldsymbol{z}')\,\mathrm{d}\boldsymbol{z}' &= \sqrt{(2\pi)^L\det(\boldsymbol{\Omega})}\int \mathcal{N}(\boldsymbol{z}-\boldsymbol{z}';\boldsymbol{0},\boldsymbol{\Omega})\,\mathcal{N}(\boldsymbol{z}';\boldsymbol{\mu}_j,\boldsymbol{P}_j)\,\mathrm{d}\boldsymbol{z}' \\
&= \sqrt{(2\pi)^L\det(\boldsymbol{\Omega})}\,\mathcal{N}(\boldsymbol{z};\boldsymbol{\mu}_j,\boldsymbol{P}_j+\boldsymbol{\Omega}),
\end{aligned} \tag{98}
$$

where we used the fact that the convolution of two normal distributions is yet another normal distribution with the summed means and covariance matrices. The outer integral of Eq. 97 is then also written as a convolution, yielding $\alpha_{ij}$:

$$\begin{aligned}\alpha_{ij} &= \int p_i(\boldsymbol{z})\left(\int a(\boldsymbol{z},\boldsymbol{z}')p_j(\boldsymbol{z}')\,\mathrm{d}\boldsymbol{z}'\right)\mathrm{d}\boldsymbol{z} \\ &= \sqrt{(2\pi)^L\det(\boldsymbol{\Omega})}\int p_i(\boldsymbol{z})\mathcal{N}(\boldsymbol{z};\boldsymbol{\mu}_j,\boldsymbol{P}_j+\boldsymbol{\Omega})\,\mathrm{d}\boldsymbol{z} \\ &= \sqrt{(2\pi)^L\det(\boldsymbol{\Omega})}\int \mathcal{N}(\boldsymbol{z};\boldsymbol{\mu}_i,\boldsymbol{P}_i)\,\mathcal{N}(\boldsymbol{z};\boldsymbol{\mu}_j,\boldsymbol{P}_j+\boldsymbol{\Omega})\,\mathrm{d}\boldsymbol{z} \\ &= \sqrt{(2\pi)^L\det(\boldsymbol{\Omega})}\int \mathcal{N}(\boldsymbol{\mu}_i-\boldsymbol{z};\boldsymbol{0},\boldsymbol{P}_i)\,\mathcal{N}(\boldsymbol{z};\boldsymbol{\mu}_j,\boldsymbol{P}_j+\boldsymbol{\Omega})\,\mathrm{d}\boldsymbol{z} \\ &= \sqrt{(2\pi)^L\det(\boldsymbol{\Omega})}\,\mathcal{N}(\boldsymbol{\mu}_i;\boldsymbol{\mu}_j,\boldsymbol{P}_i+\boldsymbol{P}_j+\boldsymbol{\Omega}) \\ &= \sqrt{\frac{\det(\boldsymbol{\Omega})}{\det(\boldsymbol{P}_i+\boldsymbol{P}_j+\boldsymbol{\Omega})}}\exp\!\left(-\frac{1}{2}(\boldsymbol{\mu}_i-\boldsymbol{\mu}_j)(\boldsymbol{P}_i+\boldsymbol{P}_j+\boldsymbol{\Omega})^{-1}(\boldsymbol{\mu}_i-\boldsymbol{\mu}_j)\right).\end{aligned} \tag{99}$$

The total width of this function is $\boldsymbol{P}_i+\boldsymbol{P}_j+\boldsymbol{\Omega}$. Within a single species therefore ($i=j$), the total width is $2\boldsymbol{P}+\boldsymbol{\Omega}$. Since $\boldsymbol{W}=\boldsymbol{\Omega}/2$ (the within-phenotype niche width is half of $\boldsymbol{\Omega}$; Section 3), the total width reads $2(\boldsymbol{P}+\boldsymbol{W})$. The covariance $2\boldsymbol{P}$ is also known as the between-phenotype niche width (Taper and Case 1985). Therefore the total width of $\alpha_{ii}$ is the between-phenotype niche width plus twice the within-phenotype niche width.

Obtaining $\boldsymbol{\beta}_{ij}$ is simply a matter of using Eq. 90 and $\boldsymbol{\beta}_{ij}=\partial\alpha_{ij}/\partial\boldsymbol{\mu}_i$ (Eq. 94):

$$\boldsymbol{\beta}_{ij}=\frac{\partial\alpha_{ij}}{\partial\boldsymbol{\mu}_i}=\sqrt{(2\pi)^L\det(\boldsymbol{\Omega})}\frac{\partial}{\partial\boldsymbol{\mu}_i}\mathcal{N}(\boldsymbol{\mu}_i;\boldsymbol{\mu}_j,\boldsymbol{P}_i+\boldsymbol{P}_j+\boldsymbol{\Omega})=(\boldsymbol{P}_i+\boldsymbol{P}_j+\boldsymbol{\Omega})^{-1}(\boldsymbol{\mu}_j-\boldsymbol{\mu}_i)\alpha_{ij}. \tag{100}$$

Similarly, we get $\boldsymbol{\Gamma}_{ij}$ from Eq. 92 and $\boldsymbol{\Gamma}_{ij}=\partial^2\alpha_{ij}/(\partial\boldsymbol{\mu}_i\circ\partial\boldsymbol{\mu}_i)$ (Eq. 96):

$$\begin{aligned}\boldsymbol{\Gamma}_{ij} &= \frac{\partial^2\alpha_{ij}}{\partial\boldsymbol{\mu}_i\circ\partial\boldsymbol{\mu}_i} = \sqrt{(2\pi)^L\det(\boldsymbol{\Omega})}\frac{\partial^2}{\partial\boldsymbol{\mu}_i\circ\partial\boldsymbol{\mu}_i}\mathcal{N}(\boldsymbol{\mu}_i;\boldsymbol{\mu}_j,\boldsymbol{P}_i+\boldsymbol{P}_j+\boldsymbol{\Omega}) \\ &= (\boldsymbol{P}_i+\boldsymbol{P}_j+\boldsymbol{\Omega})^{-1}\big[(\boldsymbol{\mu}_i-\boldsymbol{\mu}_j)\circ(\boldsymbol{\mu}_i-\boldsymbol{\mu}_j)-(\boldsymbol{P}_i+\boldsymbol{P}_j+\boldsymbol{\Omega})\big](\boldsymbol{P}_i+\boldsymbol{P}_j+\boldsymbol{\Omega})^{-1}\alpha_{ij}.\end{aligned} \tag{101}$$

We now turn to specifying $b_i$, $\boldsymbol{g}_i$, and $\boldsymbol{Q}_i$. They take on different forms depending on our choice of quadratic or quartic mortality functions (Eq. 76). We consider each in turn.

- Quadratic function: $b(\boldsymbol{z})=1-(\boldsymbol{z}\boldsymbol{z})/\theta^2$, where $(\boldsymbol{z}\boldsymbol{z})$ is again the scalar product of $\boldsymbol{z}$ with itself. We obtain $b_i$ from Eq. 80 via direct integration:

$$b_i=\int\left(1-\frac{(\boldsymbol{z}\boldsymbol{z})}{\theta^2}\right)p_i(\boldsymbol{z})\,\mathrm{d}\boldsymbol{z}=1-\frac{\mathrm{trace}(\boldsymbol{P}_i)+(\boldsymbol{\mu}_i\boldsymbol{\mu}_i)}{\theta^2}. \tag{102}$$

  From $\boldsymbol{g}_i=\partial b_i/\partial\boldsymbol{\mu}_i$ and $\boldsymbol{Q}_i=\partial^2 b_i/(\partial\boldsymbol{\mu}_i\circ\partial\boldsymbol{\mu}_i)$ (Eqs. 93 and 95):

$$\boldsymbol{g}_i=\frac{\partial b_i}{\partial\boldsymbol{\mu}_i}=-\frac{2}{\theta^2}\boldsymbol{\mu}_i,\qquad \boldsymbol{Q}_i=\frac{\partial^2 b_i}{\partial\boldsymbol{\mu}_i\circ\partial\boldsymbol{\mu}_i}=-\frac{2}{\theta^2}\boldsymbol{I}, \tag{103}$$

  where $\boldsymbol{I}$ is the identity matrix.

- Quartic function: $b(\boldsymbol{z})=1-(\boldsymbol{z}\boldsymbol{z})^2/\theta^4$. Again by direct integration,

$$\begin{aligned}b_i &= \int\left(1-\frac{(\boldsymbol{z}\boldsymbol{z})^2}{\theta^4}\right)p_i(\boldsymbol{z})\,\mathrm{d}\boldsymbol{z} \\ &= 1-\frac{\mathrm{trace}^2(\boldsymbol{P}_i)+2\mathrm{trace}(\boldsymbol{P}_i^2)+2\mathrm{trace}(\boldsymbol{P}_i)(\boldsymbol{\mu}_i\boldsymbol{\mu}_i)+4\boldsymbol{\mu}_i\boldsymbol{P}_i\boldsymbol{\mu}_i+(\boldsymbol{\mu}_i\boldsymbol{\mu}_i)^2}{\theta^4},\end{aligned} \tag{104}$$

and from $\boldsymbol{g}_i = \partial b_i / \partial \boldsymbol{\mu}_i$ and $\boldsymbol{Q}_i = \partial^2 b_i / (\partial \boldsymbol{\mu}_i \circ \partial \boldsymbol{\mu}_i)$,

$$\boldsymbol{g}_i = \frac{\partial b_i}{\partial \boldsymbol{\mu}_i} = -\frac{4}{\theta^4}[\mathrm{trace}(\boldsymbol{P}_i)\boldsymbol{\mu}_i + (\boldsymbol{\mu}_i\boldsymbol{\mu}_i)\boldsymbol{\mu}_i + 2\boldsymbol{P}_i\boldsymbol{\mu}_i], \tag{105}$$

$$\boldsymbol{Q}_i = \frac{\partial^2 b_i}{\partial \boldsymbol{\mu}_i \circ \partial \boldsymbol{\mu}_i} = -\frac{4}{\theta^4}[\mathrm{trace}(\boldsymbol{P}_i)\boldsymbol{I} + (\boldsymbol{\mu}_i\boldsymbol{\mu}_i)\boldsymbol{I} + 2(\boldsymbol{\mu}_i \circ \boldsymbol{\mu}_i) + 2\boldsymbol{P}_i]. \tag{106}$$

### 5 Calculating diversity

#### 5.1 Discrete diversity

For a discrete number of categories $C$, there is a wide range of metrics for quantifying diversity. Here we use the Hill numbers of order $q$, denoted ${}^qD$, which is a class of metrics encompassing e.g. the inverse Simpson and Shannon indices as special cases. Denoting the relative frequency of individuals in category $i$ by $f_i$, these indices read

$${}^qD = \left(\sum_{i=1}^{C} f_i^q\right)^{\frac{1}{1-q}}. \tag{107}$$

The inverse Simpson index, which is the inverse of the probability that two randomly sampled individuals belong to the same category, is recovered for $q = 2$. While the expression is singular at $q = 1$, we can interpret ${}^1D$ as the limit of Eq. 107 as $q$ approaches 1, in which case it returns the exponential of the Shannon index: ${}^1D = \exp(-\sum_i f_i \log(f_i))$ (Leinster and Cobbold 2012).

If the categories are species, then $C = S$, and these indices return the usual measures of species diversity. For discrete functional traits, equating the categories with the possible trait states will yield the community's functional diversity. For instance, in a community of lepidopteran species where individuals have either white or melanistic black wing coloration, we have $C = 2$, $f_1$ is the frequency of white individuals (irrespective of species identity), and $f_2$ the frequency of melanistic ones.

#### 5.2 Functional diversity in continuous trait spaces

For continuous traits (such as wing span, or the bill depth of darwin's finches, or the shell morphology of land snails), we are concerned with trait space coverage (Fontana et al. 2016). This means a measure of the fraction of trait space occupied by the community, and the evenness of this cover (Figure S1). There are various ways of obtaining such a measure. Here we do so by using the trait probability density (Carmona et al. 2016) of the community, and then applying standard diversity metrics to this distribution.

The community-wide trait probability density function, $\mathcal{D}(\boldsymbol{z})$, is the normalized sum of the trait distributions of all species:

$$\mathcal{D}(\boldsymbol{z}) = \frac{\sum_{i=1}^{S} N_i p_i(\boldsymbol{z})}{\int \sum_{k=1}^{S} N_k p_k(\boldsymbol{z})\,\mathrm{d}\boldsymbol{z}}. \tag{108}$$

The numerator is the sum of all one-species trait distributions (its shape given by $p_i(\boldsymbol{z})$ and the area under the curve by the population density $N_i$), while the denominator is a normalization factor, making sure that $\mathcal{D}(\boldsymbol{z})$ integrates to unity. The expression can be simplified:

$$\mathcal{D}(\boldsymbol{z}) = \frac{\sum_{i=1}^{S} N_i p_i(\boldsymbol{z})}{\sum_{k=1}^{S} N_k \underbrace{\int p_k(\boldsymbol{z})\,\mathrm{d}\boldsymbol{z}}_{1\ (\text{Eq. } 26)}} = \frac{\sum_{i=1}^{S} N_i p_i(\boldsymbol{z})}{\sum_{k=1}^{S} N_k} = \sum_{i=1}^{S} f_i p_i(\boldsymbol{z}), \tag{109}$$

where $f_i = N_i / \sum_{k=1}^{S} N_k$ is the relative frequency of species $i$ in the community. It then appears sensible to define (differential) functional diversity measures by simply replacing, in Eq. 107, $f_i$ with $\mathcal{D}(\boldsymbol{z})$ and the sum with an integral. Unfortunately, doing so loses many good properties of ${}^qD$—most problematically, it will no longer be invariant to changing the units of the traits in this naive continuum limit.

Instead, one should first divide the trait space into a mesh with grid size $\Delta$ (so the volume of each cell, in $L$ trait dimensions, is $\Delta^L$), yielding a discretization of trait space. We then evaluate $\mathcal{D}(\boldsymbol{z})$ in each cell, resulting in the value $\mathcal{D}_i$ in cell $i$. Finally, after normalizing the $\mathcal{D}_i$ with their sum to get $\hat{\mathcal{D}}_i = \mathcal{D}_i / \sum_k \mathcal{D}_k$, the functional diversity

$$ {}^qD = \left( \sum_{i=1}^{C} \hat{\mathcal{D}}^q \right)^{\frac{1}{1-q}} \tag{110} $$

is obtained as before, with $C$ now being the total number of grid cells.

To get an accurate diversity estimate, the trait space should be sufficiently finely discretized to make the difference between the $\mathcal{D}_i$ values in adjacent cells negligible—i.e., $\Delta$ should be small (Petchey and Gaston 2006). However, ${}^qD$ diverges as $\Delta \to 0$. For instance, if $\mathcal{D}(\boldsymbol{z})$ is a uniform distribution in the unit (hyper)cube of dimension $L$, then Eq. 110 gives, for an arbitrary $\Delta$,

$$ {}^qD = \left( \sum_{i=1}^{C} \Delta^{Lq} \right)^{\frac{1}{1-q}} = \Bigg( \Delta^{Lq} \underbrace{\sum_{i=1}^{C} 1}_{1/\Delta^L} \Bigg)^{\frac{1}{1-q}} = \left( \Delta^{L(q-1)} \right)^{\frac{1}{1-q}} = \Delta^{-L}. \tag{111} $$

This increases without bounds as $\Delta$ approaches zero. That, however, does not matter as long as we are interested in a relative measure of diversity across communities. Then, as long as the same region of trait space is discretized using the same $\Delta$ for all communities, and $\Delta$ is sufficiently small to yield a good resolution of trait space, the relative functional diversity values will be accurately captured. Making the community with uniform $\mathcal{D}(\boldsymbol{z})$ in the unit $L$-dimensional cube the basis for comparison, we define the functional diversity index by normalizing Eq. 110 with 111:

$$ {}^qD = \Delta^L \left( \sum_{i=1}^{C} \hat{\mathcal{D}}^q \right)^{\frac{1}{1-q}}, \tag{112} $$

which is finite and well-defined even for an arbitrarily small $\Delta$.

As an example, consider a community with a single trait ($L = 1$) that can take values between 0 and 1, and is uniform in the union of [0, 1/4] and [3/4, 1] (thus the middle half of the trait axis, [1/4, 3/4], is empty). Therefore, $\mathcal{D}(z) = 0$ for $1/4 \le z \le 3/4$ and 2 otherwise. Discretizing the [0, 1] interval with a grid size of $\Delta$, $\hat{\mathcal{D}}_i = 2\Delta$ for half the cells and $\hat{\mathcal{D}}_i = 0$ for the other (middle) half. Then, by Eq. 112, we get ${}^qD = \Delta\,(2\Delta)^{-1} = 1/2$, which is finite and meaningful even in the $\Delta \to 0$ limit.

In the main text, we always show results with $q = 2$, both for species diversity (Eq. 107) and functional diversity (Eq. 112). Here in the supplement, we indicate the value of $q$ in the figure captions.

#### 5.3 Distance-dependent functional diversity metrics

The diversity metrics of Eqs. 107 and 112 are distance-independent: only the extent and evenness of trait space coverage matter, not which parts of trait space are covered. However, there are diversity metrics which would score panel A of Figure S1 as having lower diversity than panel B, on the grounds that in B, the covered regions are further apart and therefore more varied.

This approach is indispensable when one does not have access to trait probability densities in a continuous trait space. In that case, since the volume taken up by any number of discrete points in trait space is zero, the only way to account for the fact that a set of 100 points uniformly distributed in the [0, 1] range of a single trait is more diverse than another 100 distributed in [0.5, 0.6] (say) is to use a distance-sensitive metric. One such family of metrics is the diversity of order $q$ (Leinster and Cobbold 2012), defined as

$$ {}^{q}D^{\boldsymbol{Z}} = \left( \sum_i f_i \left( \sum_j Z_{ij} f_j \right)^{q-1} \right)^{\frac{1}{1-q}} . \tag{113} $$

Here $q$ is a sensitivity parameter (adjusting the relative weight assigned to common vs. rare types), $f_i$ is the frequency of individuals having the $i$th trait value, and $\boldsymbol{Z}$ is a matrix with entries $Z_{ij}$ that depends on the distance $d_{ij}$ between the $i$th and $j$th trait values. One possible and common choice for $Z_{ij}$ is

$$ Z_{ij} = \exp\left( -\frac{d_{ij}}{\xi} \right), \tag{114} $$

where $\xi$ is a parameter controlling the distance beyond which two traits are sufficiently far apart to count as significantly different.

Since our approach uses trait probability densities and not discrete individuals, and we are interested in trait coverage in particular, it makes less sense to use distance-dependent metrics—from the point of view of the extent and evenness of the trait space covered, panels A and B of Figure S1 should count as equally diverse. Unless there are independent reasons for believing that further-away regions covered by traits are more valuable for functional diversity, this is the right approach. Nevertheless, we did repeat our analyses measuring functional diversity using Eq. 113 as well (Section 8).

### 6 Species- and functional diversity in model communities

#### 6.1 Simulation protocol

To explore the model's behavior, the following parameters were varied:

- Number of trait dimensions $L \in \{1, 2, 3\}$.
- Number of initial species $S \in \{2, 3, \dots, 25\}$.
- Initial genetic covariances $\boldsymbol{G}_i(0) \in$ {low, high}. They are generated via $\boldsymbol{G}_i(0) = \boldsymbol{U}_i \boldsymbol{B}_i \boldsymbol{U}_i^{\mathrm{T}}$, where $\boldsymbol{U}_i$ is a random orthogonal matrix and $\boldsymbol{B}_i$ is diagonal with nonzero entries sampled uniformly and independently from either [0.01, 0.05] (low) or [0.05, 0.1] (high).
- Environmental trait covariances $\boldsymbol{E}_i \in$ {low, medium, high}. $\boldsymbol{E}_i$ is a diagonal matrix with nonzero entries sampled uniformly and independently from either [0.005, 0.008] (low), [0.015, 0.018] (medium), or [0.025, 0.028] (high).
- Competition width $\omega \in$ {low, high}. We assume isotropic competition, meaning that the matrix $\boldsymbol{\Omega}$ (Eq. 72) can be written in the form $\boldsymbol{\Omega} = (\omega^2/2)\boldsymbol{I}$, where $\omega^2$ is the competition width in all directions in trait space, $\boldsymbol{I}$ is the identity matrix, and the factor 1/2 is introduced for convenience. The low and high values of $\omega$ depend on trait dimension:
  - for $L = 1$, $\omega$ is either 0.1 (low) or 0.15 (high);
  - for $L = 2$, $\omega$ is either 0.25 (low) or 0.3 (high);
  - for $L = 3$, $\omega$ is either 0.4 (low) or 0.45 (high).

- Shape of intrinsic growth function $b(z) \in \{\text{quadratic, quartic}\}$ (Eq. 76).

This leads to $3 \times 24 \times 2 \times 3 \times 2 \times 2 = 1728$ unique parameter combinations. Ten replicates were run for each parameterization. Each run was integrated for $10^{10}$ time units, to make sure the model communities reach an eco-evolutionarily stable equilibrium. Results are shown in Figures S2-S5.

A note about why environmental variances were very slightly randomized. In the model, groups of species will often undergo convergent evolution, evolving identical $\boldsymbol{\mu}_i$ and $\boldsymbol{G}_i$. Then, if all the $\boldsymbol{E}_i$ are equal as well, the species become indistinguishable, and therefore will coexist neutrally. However, this neutral coexistence is broken by the slightest difference in $\boldsymbol{E}_i$, with one of the species (possessing the "best" $\boldsymbol{E}_i$ value) driving all others in the group extinct. Since precisely equal $\boldsymbol{E}_i$ values across species are biologically implausible, this outcome would be artifactual. Slightly randomizing the $\boldsymbol{E}_i$ eliminates this artificial inflation of species diversity via neutrality.

#### 6.2 More intrinsic growth shapes in one trait dimension

To further demonstrate that the results are insensitive to the shape of $b(z)$, we implemented three additional forms of this function:

- Rectangular function:

$$b(z) = \begin{cases} 1 & \text{if } -\theta \le z \le \theta \\ 0 & \text{otherwise} \end{cases} \tag{115}$$

- Triangular function:

$$b(z) = \begin{cases} \dfrac{z+\theta}{2\theta} & \text{if } -\theta \le z \le \theta \\ 0 & \text{otherwise} \end{cases} \tag{116}$$

- Bimodal function:

$$b(z) = 2(\theta^2 - z^2)\frac{(3+\sqrt{5})z^2 + 2\theta^2}{5\theta^4} \tag{117}$$

The value of $\theta$, as before, is fixed at $\theta = 1/2$. The three functions are depicted in Figure S6. Since these functions cannot be integrated analytically for more than one dimension in Eq. 80, we restrict the analysis to single-dimensional trait spaces. Otherwise, we execute the same protocol as in Section 6.1. The results (Figures S7-S8) are as before: functional diversity declines with species diversity.

#### 6.3 Ecosystem functioning

One common measure of how well an ecosystem functions is the total community density (or biomass), $\sum_{i=1}^{S} N_i$. Another way of characterizing ecosystem functioning is through the number and quantity of resources utilized by the community as a whole. Here we show that, in our model, the two are identical up to an irrelevant factor.

To do so, we obtain the expression for the total resource quantity utilized by the community, $F$. Since Eq. 62 gives the available resource level as its maximum density $R_0(\boldsymbol{y})$ minus what has been consumed, $\sum_{j=1}^{S} \int u(\boldsymbol{z}', \boldsymbol{y}) N_j p_j(\boldsymbol{z}')\, \mathrm{d}\boldsymbol{z}'$, this second term measures how much of resource $R(\boldsymbol{y})$ is being utilized. So the total resource use $F$ is the integral of this quantity with respect to $\boldsymbol{y}$:

$$F = \int \left( \sum_{j=1}^{S} \int u(\boldsymbol{z}, \boldsymbol{y}) N_j p_j(\boldsymbol{z})\, \mathrm{d}\boldsymbol{z} \right) \mathrm{d}\boldsymbol{y}. \tag{118}$$

Substituting $u(\boldsymbol{z}, \boldsymbol{y})$ from Eq. 69 and $p_j(\boldsymbol{z})$ from Eq. 24, and rearranging:

$$F = \sum_{j=1}^{S} N_j \left[ (2\pi)^L \det(\boldsymbol{\Omega}) \right]^{1/4} \int \left( \int \mathcal{N}(\boldsymbol{z}; \boldsymbol{y}, \boldsymbol{\Omega}/2) \mathcal{N}(\boldsymbol{z}; \boldsymbol{\mu}_j, \boldsymbol{P}_j) \, \mathrm{d}\boldsymbol{z} \right) \mathrm{d}\boldsymbol{y}. \tag{119}$$

The parenthesized integral is the convolution of two normal functions, which is itself normal with the summed means and covariance matrices:

$$F = \sum_{j=1}^{S} N_j \left[ (2\pi)^L \det(\boldsymbol{\Omega}) \right]^{1/4} \int \mathcal{N}(\boldsymbol{\mu}_j; \boldsymbol{y}, \boldsymbol{\Omega}/2 + \boldsymbol{P}_j) \, \mathrm{d}\boldsymbol{y}. \tag{120}$$

The integral over a normal distribution is 1, and the factor multiplying $N_j$ is a constant that can be brought out before the summation. Therefore, we end up with

$$F = \left[ (2\pi)^L \det(\boldsymbol{\Omega}) \right]^{1/4} \sum_{j=1}^{S} N_j, \tag{121}$$

which is indeed the total community density times a constant.

Figure S9 shows the behavior of total community density, $\sum_{i=1}^{S} N_i$, as species diversity increases. We find that ecosystem functioning is invariably enhanced by species diversity. This is despite the fact that functional trait diversity actually diminishes. Thus, species diversity is beneficial for ecosystem functioning—but crucially, not because functional diversity increases with species diversity.

### 7 Species- and functional diversity in the empirical snail communities

#### 7.1 General methodology

Empirical data are organized in tables where each row is a single individual, and columns record individuals' species identity, the community they belong to, and their morphological trait measurements. To obtain species diversity, we calculate the frequencies $f_i$ in each community as the number of individuals of species $i$ divided by the total number of individuals in that community. We then apply the diversity index of Eq. 107, with $C$ equal to the community-specific number of species. To obtain functional diversity, we first standardize every morphological variable by subtracting the mean and dividing the result by the standard deviation, calculated across the whole dataset. Then we decide the subset of morphological variables to be considered, making up the trait vector $\boldsymbol{z}$.

Next, we construct the community-wide trait probability density function $\mathcal{D}(\boldsymbol{z})$ from Eq. 109, using the frequencies $f_i$ and the fitted species-specific trait distributions $p_i(\boldsymbol{z})$. This fitting was done via one of two methods. First, multivariate normal distributions were fitted using the unbiased maximum likelihood estimators for the mean $\hat{\boldsymbol{\mu}}_i$ and covariance matrix $\hat{\boldsymbol{P}}_i$ of species $i$:

$$\hat{\boldsymbol{\mu}}_i = \frac{1}{n_i} \sum_{k=1}^{n_i} \boldsymbol{x}_{i,k} \tag{122}$$

and

$$\hat{\boldsymbol{P}}_i = \frac{1}{n_i - 1} \sum_{k=1}^{n_i} (\boldsymbol{x}_{i,k} - \hat{\boldsymbol{\mu}}_i) \circ (\boldsymbol{x}_{i,k} - \hat{\boldsymbol{\mu}}_i), \tag{123}$$

where $n_i$ is the number of individuals of species $i$ in the given community, and $\boldsymbol{x}_{i,k}$ is the trait vector of species $i$'s individual $k$. Second, instead of assuming a multinormal shape for the trait distributions, we also approximated them using multivariate kernel smoothing (Carmona et al. 2016), obtaining a kernel density estimate from the data. This was done using the R package `ks` (R Core Team 2019).

Once $\mathcal{D}(\boldsymbol{z})$ of the community is obtained from $f_i$, $p_i(\boldsymbol{z})$, and Eq. 109, we apply Eq. 112 for the functional diversity. Discretization is done with a grid size $\Delta$ within a cubic volume of trait space. Both $\Delta$ and the side length of the cube are chosen so that subsequent decreases in $\Delta$ or increases in cube size no longer affect the relative functional diversity values across species and communities.

#### 7.2 The Galápagos land snail data

Individuals of the land snail data belong to a particular island, and within the island, they either inhabit the arid or humid vegetation zone. The distribution ranges of snail species never overlap across the arid and humid zones, so their species do not have the opportunity to interact with one another. This means that the species compositions of the humid and arid zones form effectively separate communities, and are treated here as such.

Three small satellite islands were removed from the data (CH, ED, and GA). In addition, there were only two sampled individuals of the species *Naesiotus achatinellus*; those were also removed (all other species had at least 13 individuals sampled, with most having at least 20 specimens). Snails were placed and their functional diversity evaluated in a two-dimensional trait space whose axes correspond to centroid size and the first PC axis of shell shape (Parent and Crespi 2009). This principal axis corresponds to whether shells are long and thin or compact and wide, and explains over 80% of all shell shape variation. The distribution of individuals in each subcommunity reveals that species segregate in this two-dimensional trait space, even though this segregation would not be evident by looking at only one of the two traits at a time (Figure S10).

Raw species- and functional diversity data are shown in Figure S11. This plot does not take into account the fact that islands vary greatly in their plant diversity. This is important because snail species partition the habitat according to the opportunities plants provide (leaves, trunks, logs, twigs, etc.), and it is thought that plant diversity is therefore tightly associated with habitat availability for snails. To correct for this, it is more reasonable to plot species diversity per host plant species in the given community. This is the figure shown in the main text.

The main text displays results assuming $q = 2$ in Eq. 112. We also show that the obtained diversity patterns are robust to the choice of $q$ (Figure S12).

There is one additional complication: the sampling methodology was such that relative abundances in the data do not express relative abundances on the islands. While knowing relative abundances is essential for diversity calculations, in our case the results are robust to simply randomizing abundances. We assigned a random abundance to each species, between 1 and 100, and recalculated both functional diversity and species diversity per host plant species. We then performed a linear regression and recorded whether its slope was negative. Repeating this procedure 1000 times, we found that the regression slope was negative in 911 cases. This reinforces the impression that there is no positive relationship, and even evidence of a negative one, between species- and functional diversity in this dataset.

These results were all obtained by fitting species' trait distributions with multivariate normal distributions. However, they are qualitatively unchanged if, instead of assuming that Gaussian distributions fit the data well, we instead approximate the trait distributions via kernel density estimation (Carmona et al. 2016). See Figure S13 for the results.

### 8 Distance-dependent functional diversity metrics

Figure S14 shows the same diversity information as Figure S2, except with functional diversity measured by the distance-dependent metric of Eq. 113. We set the sensitivity parameter to $q = 2$, in which case ${}^{q}D^{\mathbf{Z}}$ collapses to Rao's quadratic entropy measure (Leinster and Cobbold 2012). Figure S15 summarizes results for the empirical data. All results are qualitatively identical to those of the main text, for a wide range of distance parameters $\xi$. Switching to a distance-dependent functional diversity metric therefore does not affect our findings.

| Symbol | Description |
|---|---|
| $S$ | Number of species |
| $L$ | Dimensionality of trait space |
| $\boldsymbol{z}$ | Vector in trait space |
| $\boldsymbol{m}$ / $\boldsymbol{f}$ | Maternal / paternal genetic contribution to phenotype |
| $\boldsymbol{e}$ | Environmental contribution to phenotype |
| $\boldsymbol{p}$ | Phenotype vector of an individual; $\boldsymbol{p} = \boldsymbol{m} + \boldsymbol{f} + \boldsymbol{e}$ |
| $\boldsymbol{G}$ | Genetic covariance matrix; $\boldsymbol{G} = 2\text{cov}(\boldsymbol{m}) = 2\text{cov}(\boldsymbol{f})$ |
| $\boldsymbol{E}$ | Environmental covariance matrix; $\boldsymbol{E} = \text{cov}(\boldsymbol{e})$ |
| $\boldsymbol{P}$ | Total phenotypic covariance matrix; $\boldsymbol{P} = \boldsymbol{G} + \boldsymbol{E}$ |
| $\boldsymbol{p}_p$ / $\boldsymbol{p}_m$ | Parent / Midparent phenotype vector |
| $\boldsymbol{\mu}$ | Mean phenotype vector of species |
| $\mathcal{N}(\boldsymbol{z}; \boldsymbol{\mu}, \boldsymbol{P})$ | Multivariate normal distribution with mean $\boldsymbol{\mu}$ and covariance matrix $\boldsymbol{P}$ |
| $\boldsymbol{\mu}_{p_m,p}$ | Mean of midparent-offspring regression |
| $\boldsymbol{\Sigma}_{p_m,p}$ | Covariance matrix of midparent-offspring regression |
| $p_i(z)$ | Phenotype distribution of species $i$ (multinormal with mean $\boldsymbol{\mu_i}$ and cov $\boldsymbol{P}_i$) |
| $N_i$ | Population density of species $i$ |
| $W_i(\boldsymbol{z})$ | Discrete-time absolute fitness of species $i$'s phenotype $\boldsymbol{z}$ |
| $r_i(\boldsymbol{z})$ | Continuous-time per capita growth rate species $i$'s phenotype $\boldsymbol{z}$ |
| $\boldsymbol{z} \circ \boldsymbol{\mu}$ | Outer product of two vectors $\boldsymbol{z}$ and $\boldsymbol{\mu}$ (such that $[\boldsymbol{z} \circ \boldsymbol{\mu}]^{kl} = z^k \mu^l$) |
| $R(\boldsymbol{y})$ | Resource density of type $\boldsymbol{y}$ |
| $R_0(\boldsymbol{y})$ | Saturation density of resource type $\boldsymbol{y}$ |
| $m(\boldsymbol{z})$ | Intrinsic mortality rate of phenotype $\boldsymbol{z}$ |
| $\theta$ | Width of intrinsic mortality function; always set to $\theta = 1/2$ |
| $u(\boldsymbol{z}, \boldsymbol{y})$ | Utilization of resource $\boldsymbol{y}$ by a consumer of phenotype $\boldsymbol{z}$ |
| $b(\boldsymbol{z})$ | Effective intrinsic growth rate of phenotype $\boldsymbol{z}$ |
| $a(\boldsymbol{z}, \boldsymbol{z}')$ | Effective competition kernel between phenotypes $\boldsymbol{z}$ and $\boldsymbol{z}'$ |
| $\boldsymbol{W}$ | Breadth of resource utilization curve $u(\boldsymbol{z}, \boldsymbol{y})$ (within-phenotype niche width) |
| $\boldsymbol{\Omega}$ | Breadth of effective competition kernel $a(\boldsymbol{z}, \boldsymbol{z}')$; $\boldsymbol{\Omega} = 2\boldsymbol{W}$ |
| $\boldsymbol{I}$ | Identity matrix |
| $\omega$ | Competition width, defining $\boldsymbol{\Omega} = (\omega^2/2)\boldsymbol{I}$ |
| $b_i$ | Species-level intrinsic growth rate, for species $i$ |
| $\alpha_{ij}$ | Competition coefficient between species $i$ and $j$ |
| $\boldsymbol{B}$ | Breadth of $\alpha_{ii}$ (between-phenotype niche width); $\boldsymbol{B} = 2(\boldsymbol{P} + \boldsymbol{W})$ |
| $\boldsymbol{g}_i$ | Selection pressure on species $i$'s trait mean from growth |
| $\boldsymbol{\beta}_{ij}$ | Selection pressure on species $i$'s trait mean from competition with species $j$ |
| $\boldsymbol{Q}_i$ | Selection pressure on species $i$'s trait covariance from growth |
| $\boldsymbol{\Gamma}_{ij}$ | Selection pressure on species $i$'s trait cov from competition with species $j$ |
| $(\boldsymbol{z}\boldsymbol{z})$ | Scalar product of $\boldsymbol{z}$ with itself (equal to squared length of $\boldsymbol{z}$) |
| $\text{trace}(\boldsymbol{P})$ | Trace (sum of diagonal entries) of the matrix $\boldsymbol{P}$ |
| ${}^qD$ | Hill number (diversity index) of order $q$ |
| ${}^qD^{\boldsymbol{Z}}$ | Distance-dependent diversity index of order $q$, with similarity matrix $\boldsymbol{Z}$ |
| $\mathcal{D}(\boldsymbol{z})$ | Trait probability density, as a function of trait value $\boldsymbol{z}$ |
| $\Delta$ | Grid size of discretized trait space |
| $\hat{\mathcal{D}}_i$ | Normalized trait probability density in grid cell $i$ |
| $F$ | Total amount of resources utilized by the community |
| $\hat{\boldsymbol{\mu}}_i$ | Maximum likelihood estimate of species $i$'s trait mean |
| $\hat{\boldsymbol{P}}_i$ | Maximum likelihood estimate of species $i$'s trait covariance matrix |

**Table S1:** Table of symbols

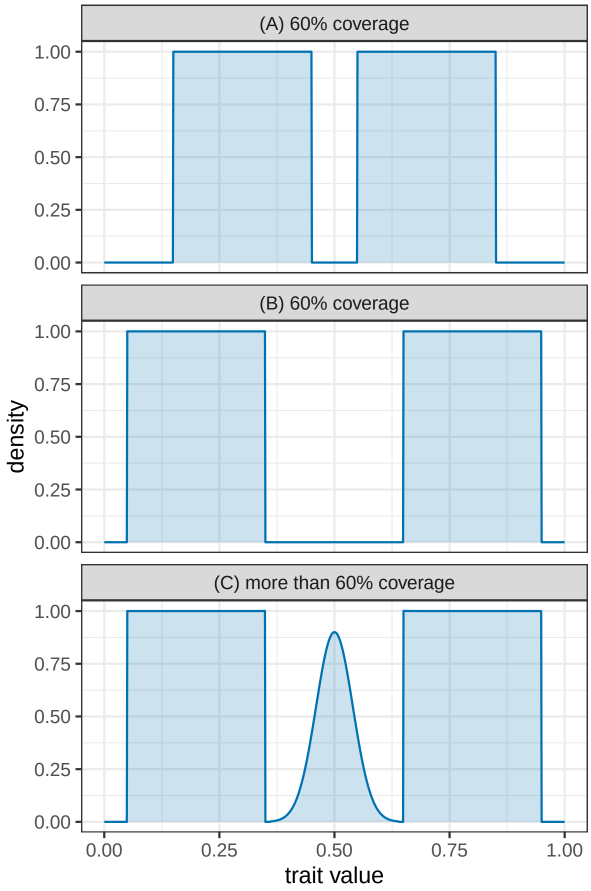


**Figure S1:** Trait probability densities of three hypothetical communities, over a one-dimensional trait space where trait values may range from 0 to 1. (A): The trait probability density has two blocks in which trait values are uniformly distributed. Everywhere else, it is zero. Thus, in total, 60% of the trait axis is uniformly covered by the trait probability density function. (B): This community is as before, except there is a larger gap between the two regions of positive density. This, however, does not change the fact that still 60% of the trait space is covered, so this community is equally functionally diverse to the previous one. (C): As (B), but with an extra positive trait region in the middle. Although the total trait space coverage is 90% (all parts are covered except for the small 5% region at each end of the trait axis), coverage is not even. To determine the functional diversity of this community, a higher-order metric is needed.

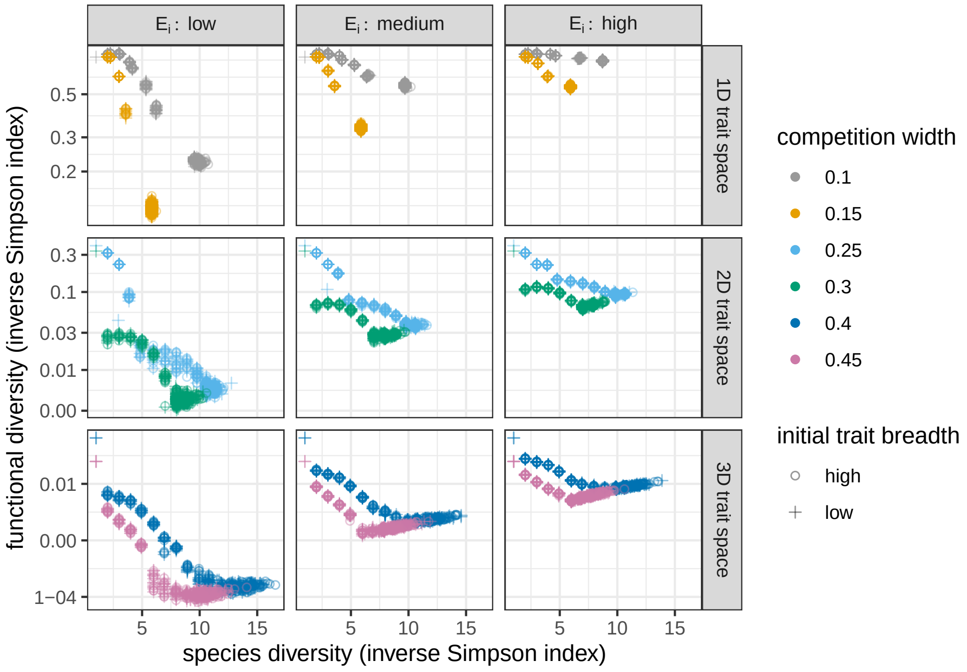


**Figure S2:** Functional diversity against species diversity, for various dimensions of trait space (rows) and levels of environmental trait variation (columns). Both diversities are measured via the inverse Simpson index ($q = 2$ in Eqs. 107 and 112), with each trait axis discretized in the range $[-1, 1]$ with $\Delta = 1/100$ for all calculations of functional diversity (Section 5). Points show results across the parameter combinations and replicates. Colors indicate different values of the competition width $\omega$; shapes correspond to results with high ($\circ$) and low ($+$) initial genetic variation. Intrinsic growth rates are quadratic (Eq. 76) for all results shown.

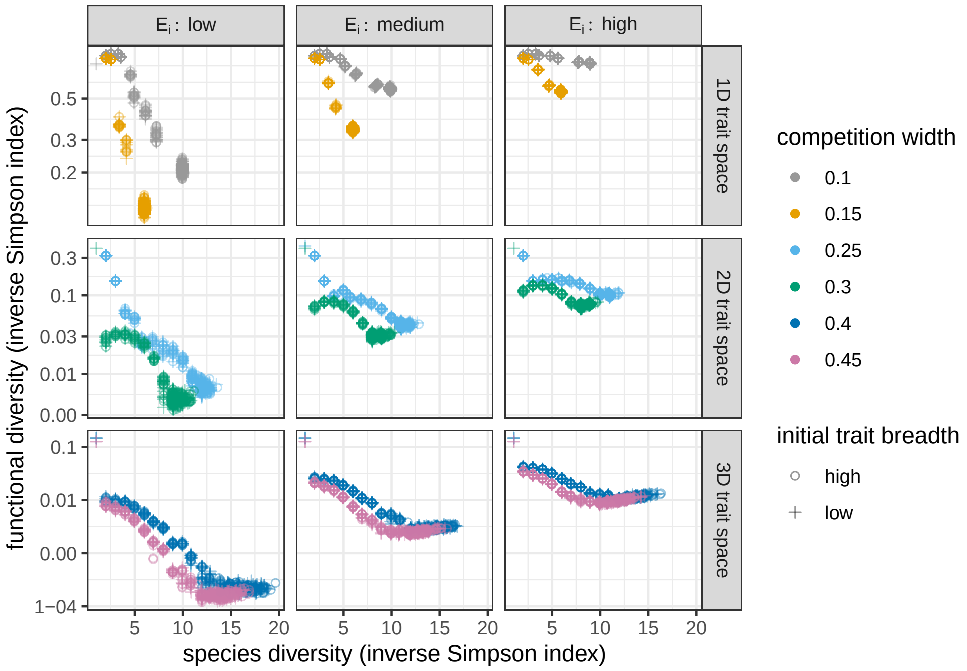


**Figure S3:** As Figure S2, with both species- and functional diversity measured by the inverse Simpson index ($q$ = 2 in Eqs. 107 and 112), and with quartic intrinsic growth rates (Eq. 76).

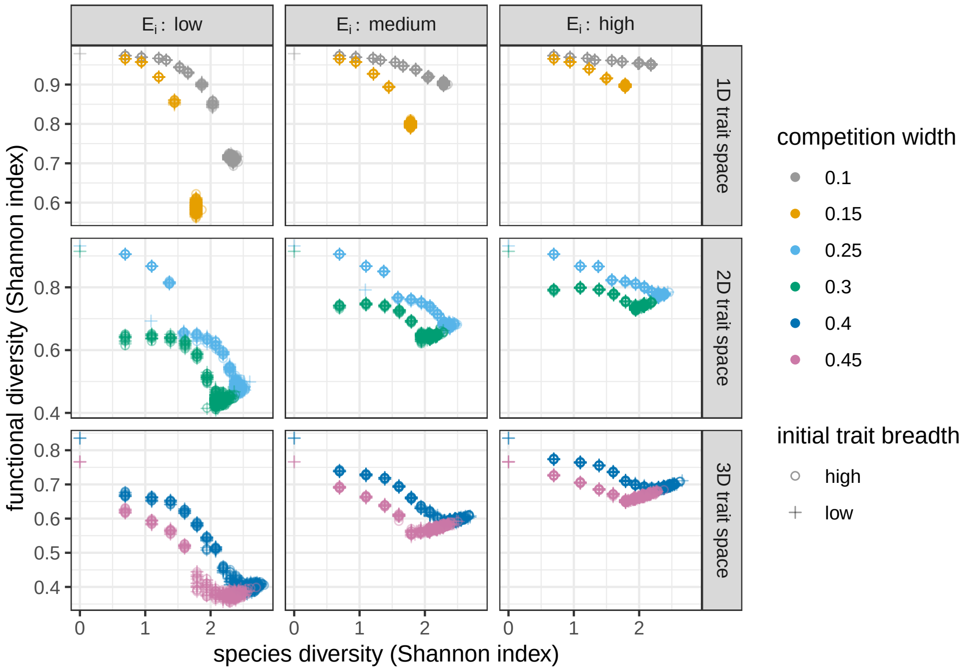


**Figure S4:** As Figure S2, with both species- and functional diversity measured by the Shannon index $\log({}^qD)$, with $q = 1$ and ${}^qD$ given by Eqs. 107 and 112, and with quadratic intrinsic growth rates (Eq. 76).

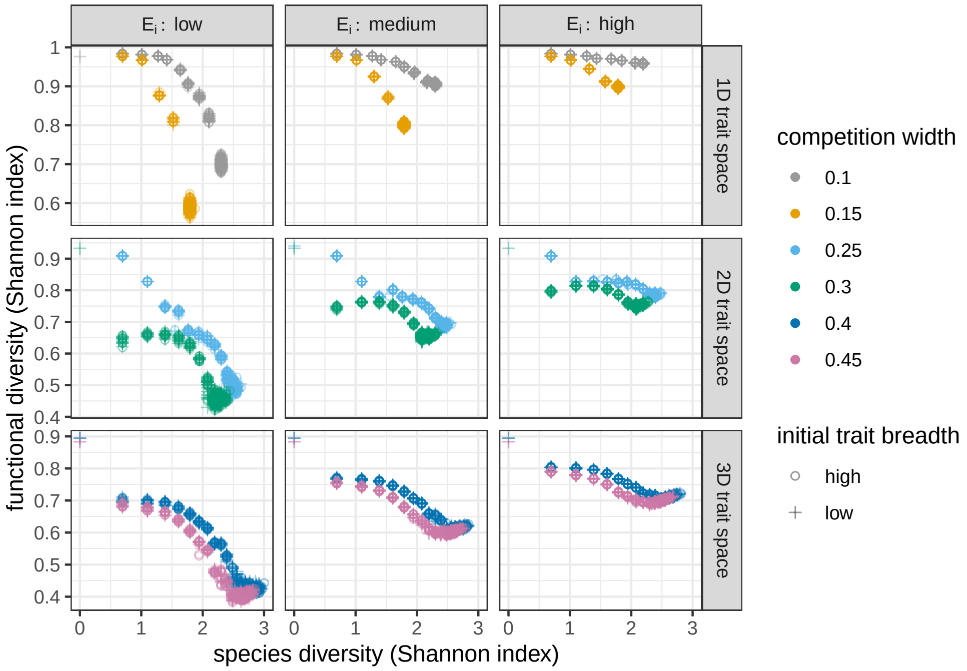


**Figure S5:** As Figure S2, with both species- and functional diversity measured by the Shannon index $\log({}^qD)$, with $q = 1$ and ${}^qD$ given by Eqs. 107 and 112, and with quartic intrinsic growth rates (Eq. 76).

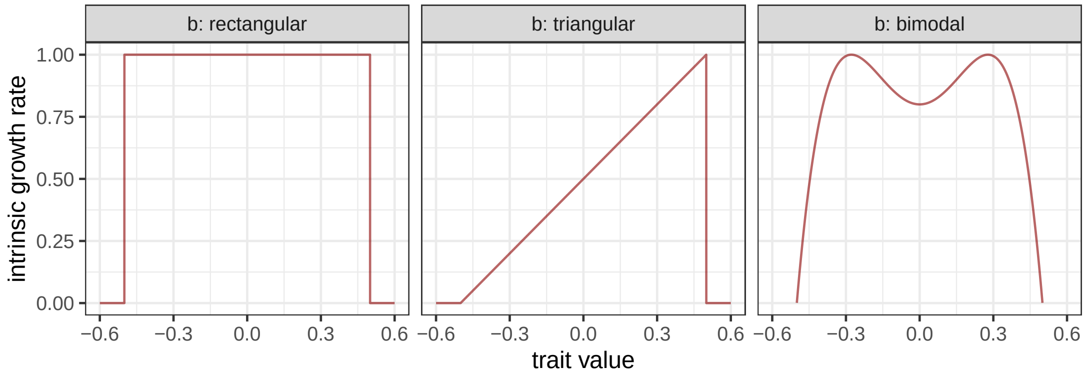


**Figure S6:** Alternative shapes for the intrinsic growth function $b(z)$: rectangular (left; Eq. 115), triangular (middle; Eq. 116), and bimodal (right; Eq. 117) forms.

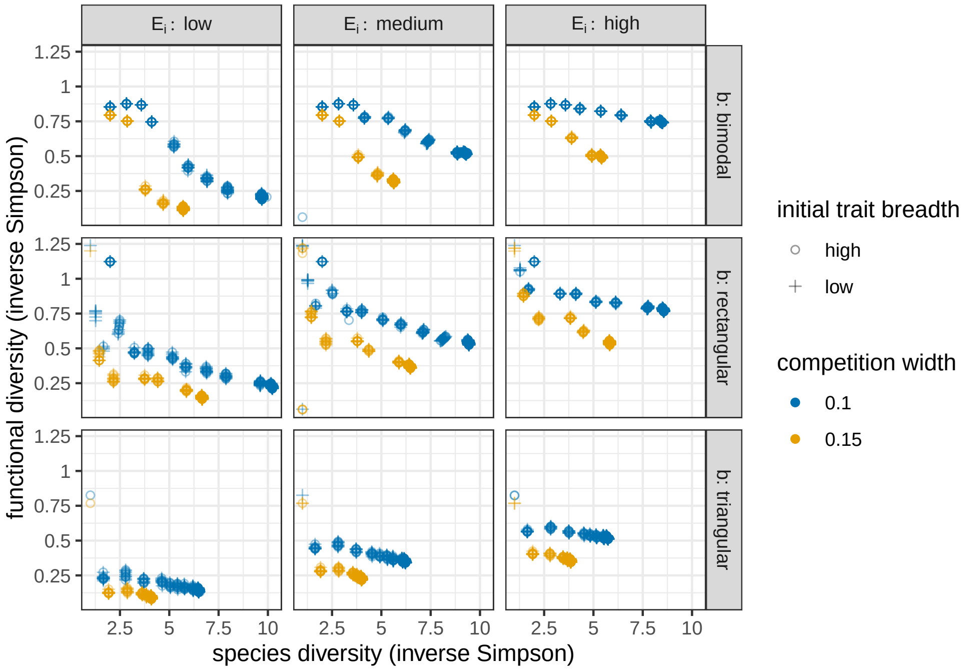


**Figure S7:** Functional diversity against species diversity, in one-dimensional trait spaces, for various shapes of the intrinsic growth function $b(z)$ (rows) and levels of environmental trait variation (columns). Both diversities are measured via the inverse Simpson index ($q = 2$ in Eqs. 107 and 112). Otherwise the figure is as Figure S4.

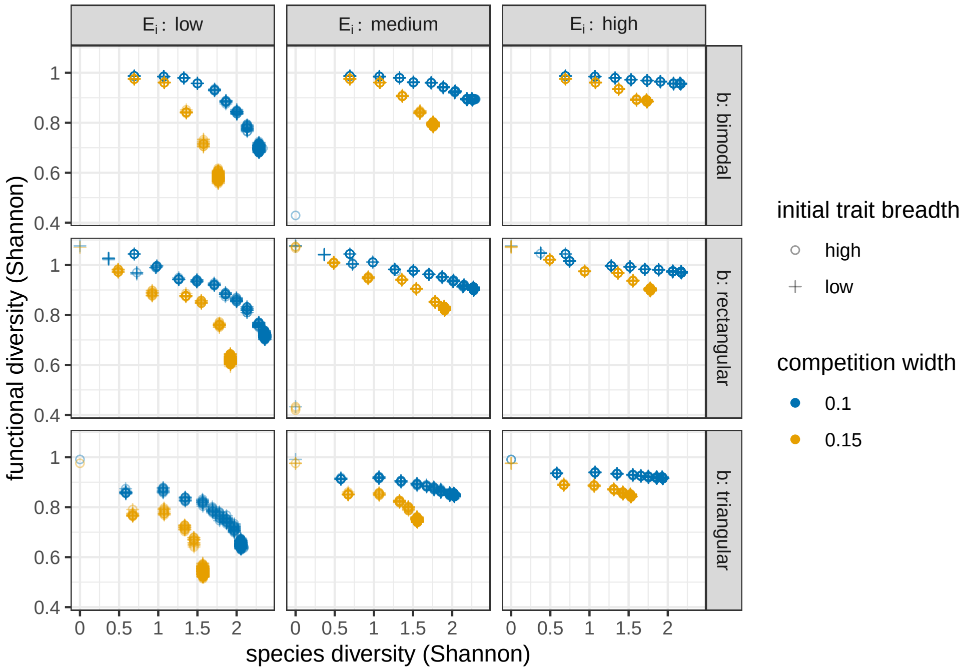


**Figure S8:** As Figure S7, but with both species- and functional diversity measured by the Shannon index $\log({}^qD)$, with $q = 1$ and ${}^qD$ given by Eqs. 107 and 112.

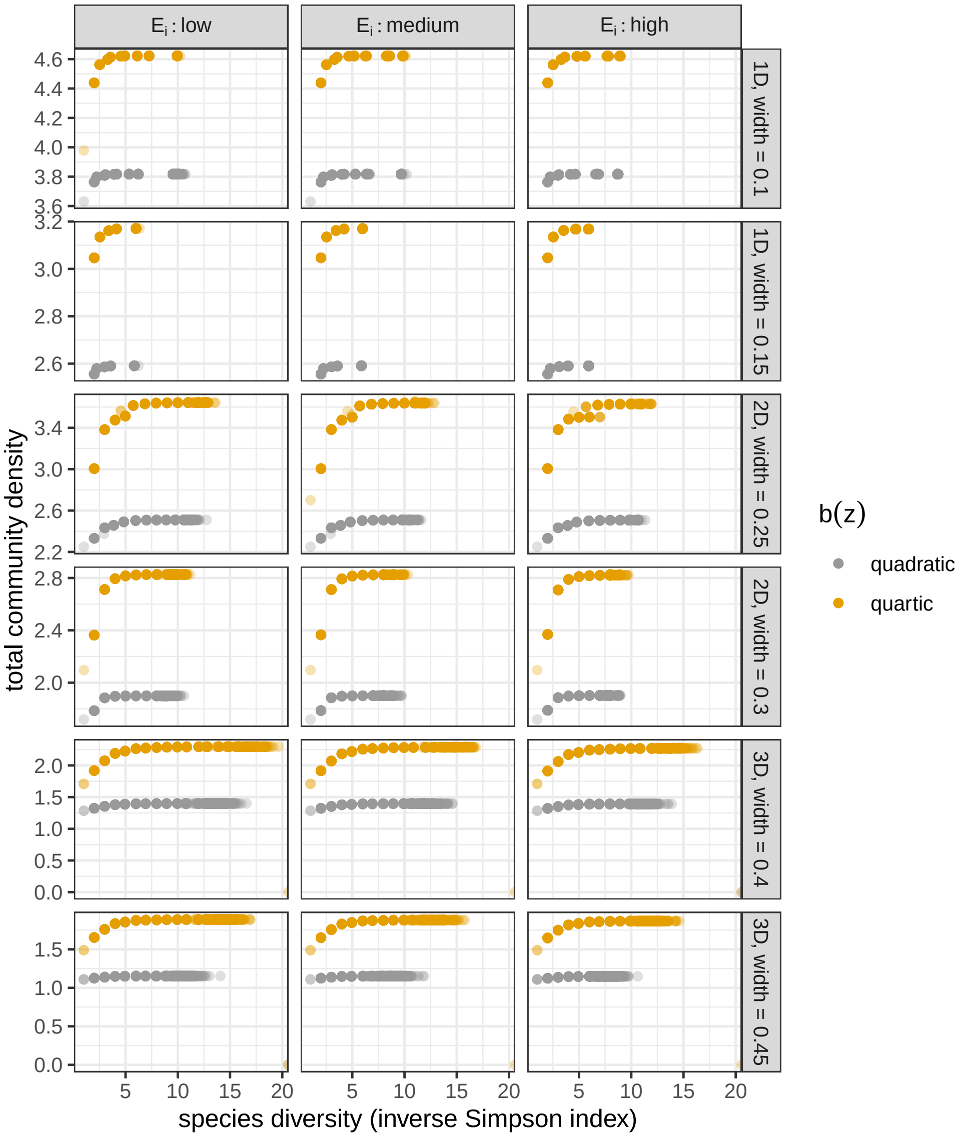


**Figure S9:** Total community density as a function of species diversity, for various vaues of trait space dimensionality and competition width (rows), levels of environmental trait variation (columns), and shapes of the intrinsic growth function $b(z)$ (colors). Regardless of these parameters, total community density always increases with increasing species diversity, indicating better ecosystem functioning. This is despite the fact that functional diversity itself declines with increasing species diversity.

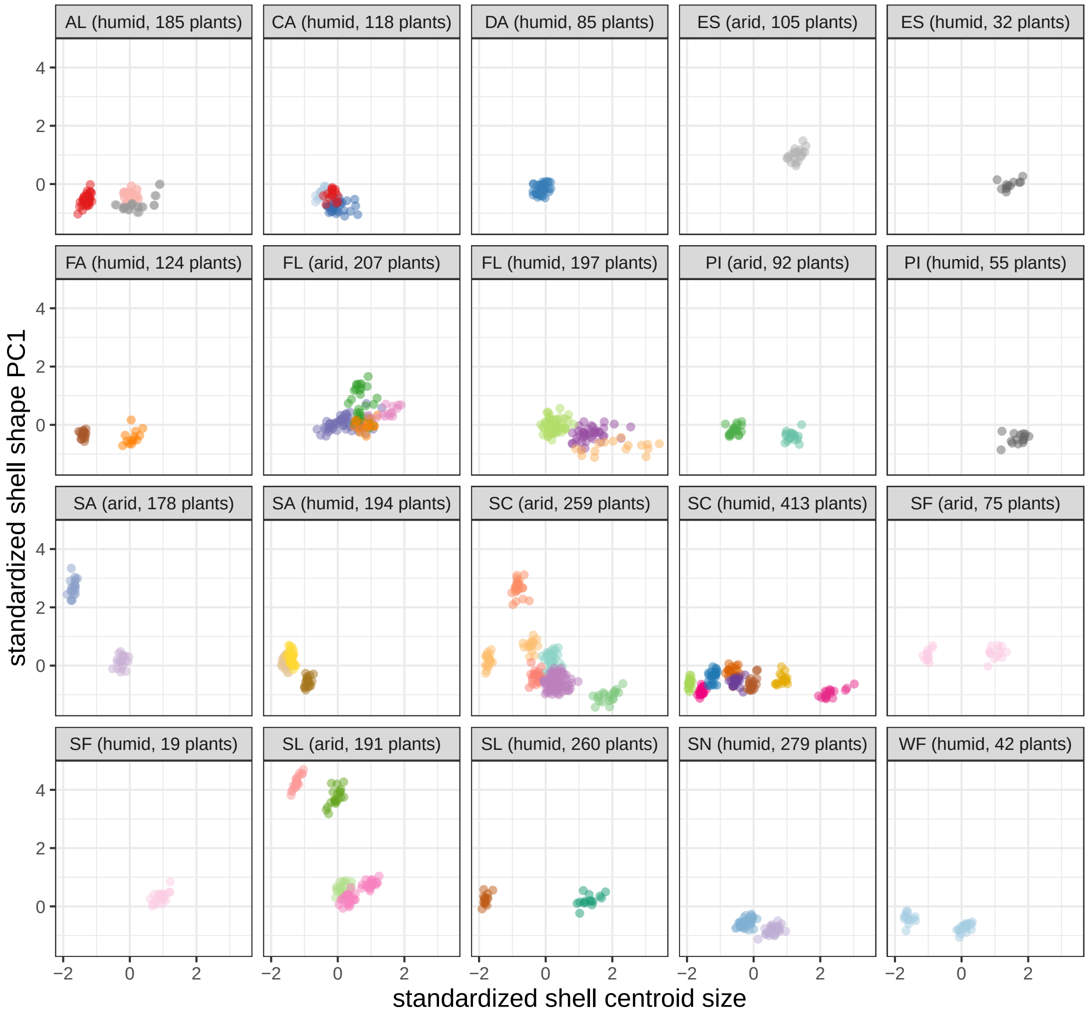


**Figure S10:** Distribution of individuals in the two-dimensional trait space spanned by the standardized shell centroid size (abscissa) and standardized shell shape (ordinate; measured by the first PC axis of shape, explaining over 80% of shape variation). Each point is an individual belonging to a species (colors). Panels show independent subcommunities, indexed by island label and habitat type within the island (arid/humid). Island name abbreviations are: Alcedo Volcano (AL), Cerro Azul Volcano (CA), Darwin Volcano (DA), Espanola (ES), Fernandina (FA), Floreana (FL), Pinzon (PI), Santiago (SA), Santa Cruz (SC), Santa Fe (SF), San Cristobal (SL), Sierra Negra Volcano (SN), Wolf Volcano (WF). Panels also indicate the number of host plant species in each community, since the number of ecological opportunities are thought to scale with this number (Parent and Crespi 2009).

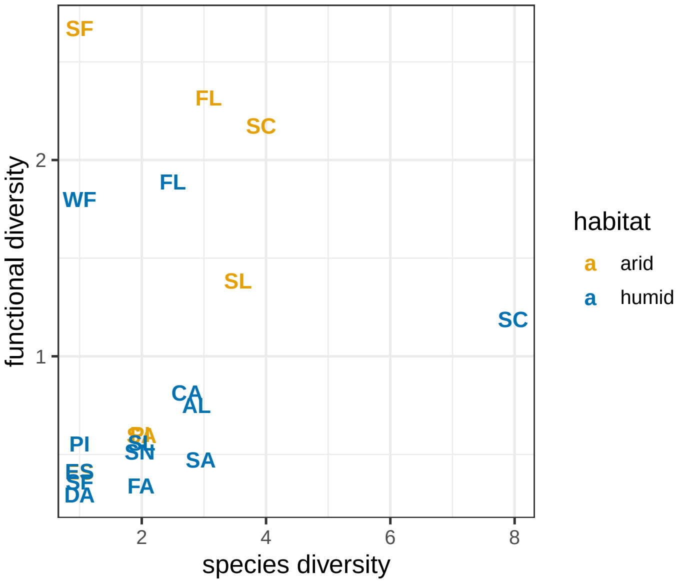


**Figure S11:** Functional diversity against species diversity, measured by the inverse Simpson index ($q = 2$ in Eqs. 107 and 112), for land snail communities on the Galápagos Islands. Species diversity is not corrected for the available host plant diversity in each community. Labels are island name abbreviations: Alcedo Volcano (AL), Cerro Azul Volcano (CA), Darwin Volcano (DA), Espanola (ES), Fernandina (FA), Floreana (FL), Pinzon (PI), Santiago (SA), Santa Cruz (SC), Santa Fe (SF), San Cristobal (SL), Sierra Negra Volcano (SN), Wolf Volcano (WF). Colors show communities in the arid (red) and humid (blue) zones of the islands, which form independent subcommunities.

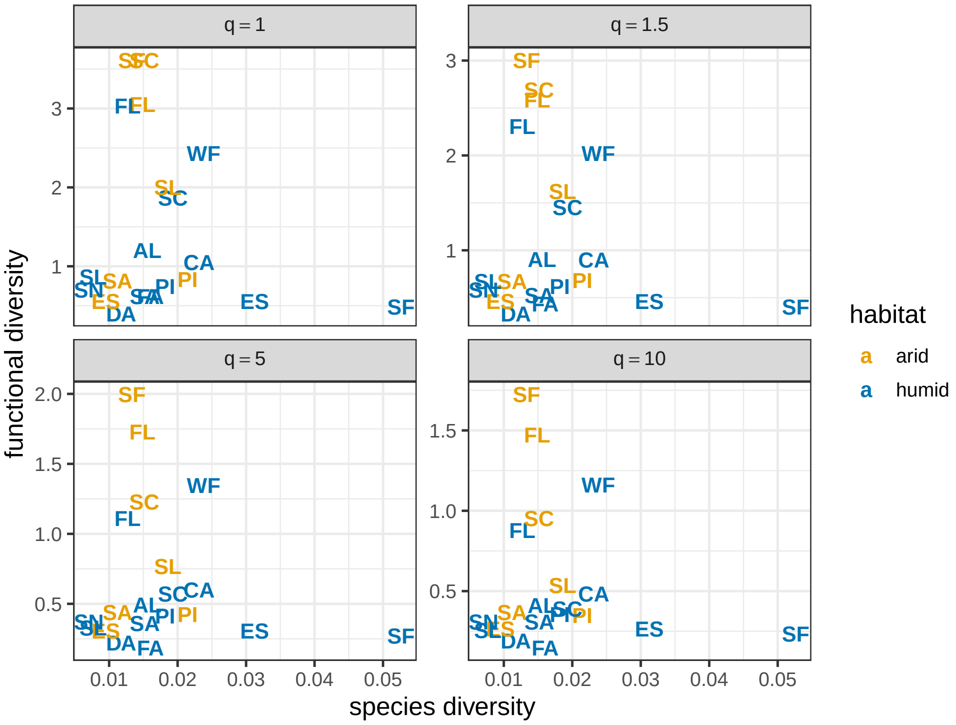


**Figure S12:** Functional diversity against species diversity for land snail communities on the Galápagos Islands. Labels are island name abbreviations: Alcedo Volcano (AL), Cerro Azul Volcano (CA), Darwin Volcano (DA), Espanola (ES), Fernandina (FA), Floreana (FL), Pinzon (PI), Santiago (SA), Santa Cruz (SC), Santa Fe (SF), San Cristobal (SL), Sierra Negra Volcano (SN), Wolf Volcano (WF). Colors show communities in the arid (red) and humid (blue) zones of the islands, which form independent subcommunities. Species diversity is measured by the inverse Simpson index ($q = 2$ in Eq. 107), while functional diversity is obtained from Eq. 112 with various values of $q$ (panels). As seen, results are robust to varying the sensitivity parameter $q$.

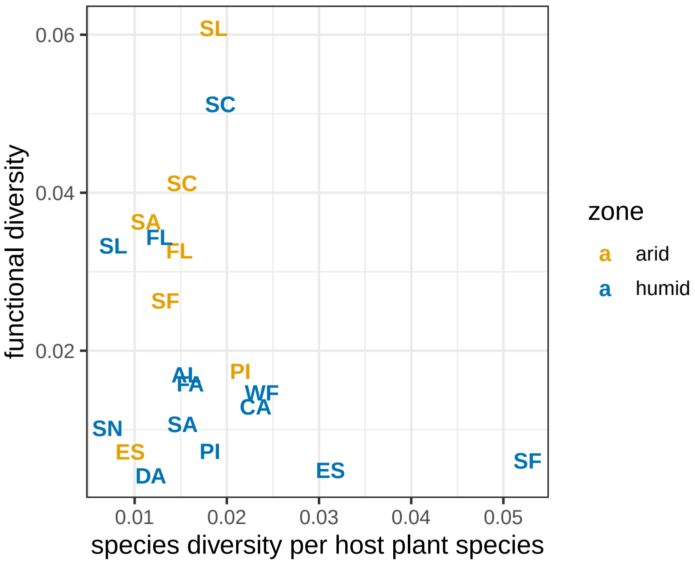


**Figure S13:** As Figure S11, but with species diversity corrected for the number of available host plants (as in the main text), and species' trait distributions estimated via kernel smoothing instead of the fitting of bivariate normal distributions. Results are qualitatively unchanged: there is still a weak negative trend which is robust to randomizing relative abundances (out of 1000 randomizations, 994 ended up with a negative slope when regressing functional diversity against species diversity).

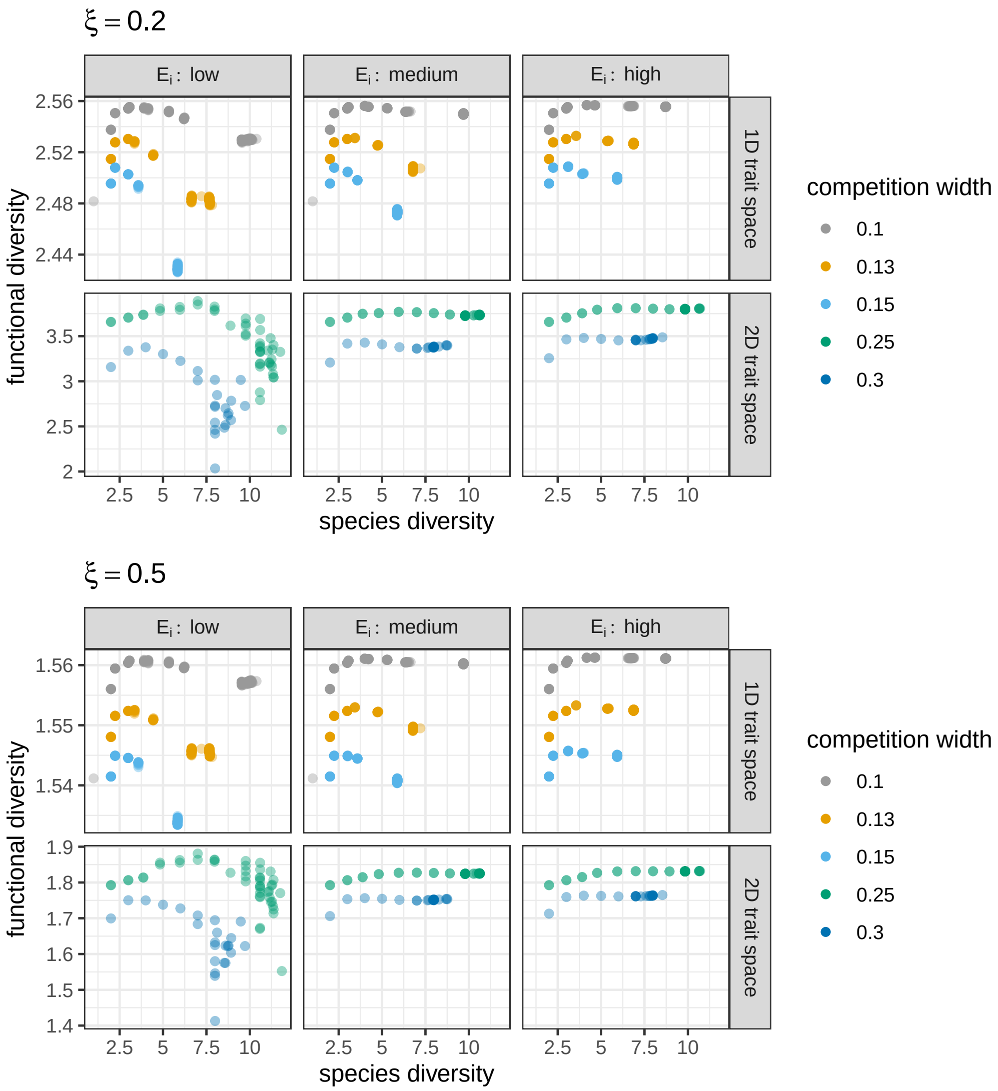


**Figure S14:** Functional diversity (measured by the distance-dependent Eq. 113 with $q = 2$) against species diversity (measured by Eq. 107 with $q = 2$), in model communities, for distance parameter $\xi = 0.2$ (top panels) and $\xi = 0.5$ (bottom panels; the effective length of each trait dimension is 1 in the model). Results are qualitatively the same as with a distance-independent functional diversity metric (Figure S2), but the increase of functional diversity at low species diversity values is more pronounced. Due to the computational intensity of obtaining this metric, we omitted three-dimensional trait spaces, and applied a somewhat crude grid resolution in the two-dimensional case (with $\Delta = 0.03$, as opposed to $\Delta = 0.01$ in one dimension).

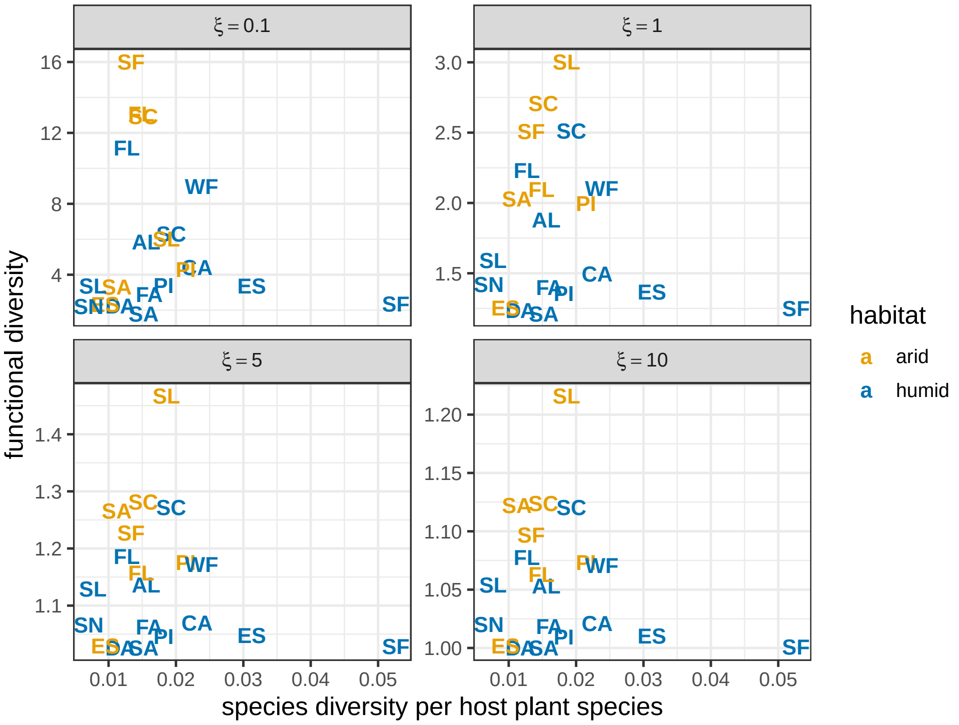


**Figure S15:** Galápagos land snail functional diversity (measured by Eq. 113 with $q = 2$) against species diversity (Eq. 107, $q = 2$). Panels show results for a wide range of $\xi$ values. In all cases, the same qualitative results are retained as with the distance-independent functional diversity indices. Since the land snail dataset has the problem that the number of sampled individuals per species is not proportional to their relative abundances, we performed the same randomization procedure as in Section 7.2 to make sure the results are robust to altering these relative abundances. Out of 100 randomizations, at least 90% had negative regression slopes for each of the four $\xi$ values, indicating that the negative relationship between species- and functional diversity is robust.
